# STING Promotes Breast Cancer Cell Survival by an Inflammatory-Independent Nuclear Pathway Enhancing the DNA Damage Response

**DOI:** 10.1101/2020.07.11.196790

**Authors:** Laura Cheradame, Ida Chiara Guerrera, Julie Gaston, Alain Schmitt, Vincent Jung, Marion Pouillard, Nina Radosevic-Robin, Mauro Modesti, Jean-Gabriel Judde, Stefano Cairo, Vincent Goffin

## Abstract

STING (Stimulator of Interferon Genes) is a well-known endoplasmic reticulum-anchored adaptor of the innate immunity that triggers the expression of inflammatory cytokines in response to pathogen infection. In cancer cells, this pro-inflammatory pathway can be activated by genomic DNA damage potentiating antitumor immune responses. Here we report that STING promotes cancer cell survival and resistance to genotoxic treatment in a cell-autonomous manner. Mechanistically, we show that STING partly localizes at the inner nuclear membrane in various breast cancer cell lines and clinical tumor samples, and interacts with several proteins of the DNA damage response (DDR). STING overexpression enhances the amount of chromatin-bound DNA-dependent Protein Kinase (DNA-PK) complex, while STING silencing impairs DDR foci formation and DNA repair efficacy. Importantly, this function of STING is independent of its canonical pro-inflammatory pathway. This study highlights a previously unappreciated cell-autonomous tumor-promoting mechanism of STING that opposes its well-documented role in tumor immunosurveillance.

**Graphical abstract:** 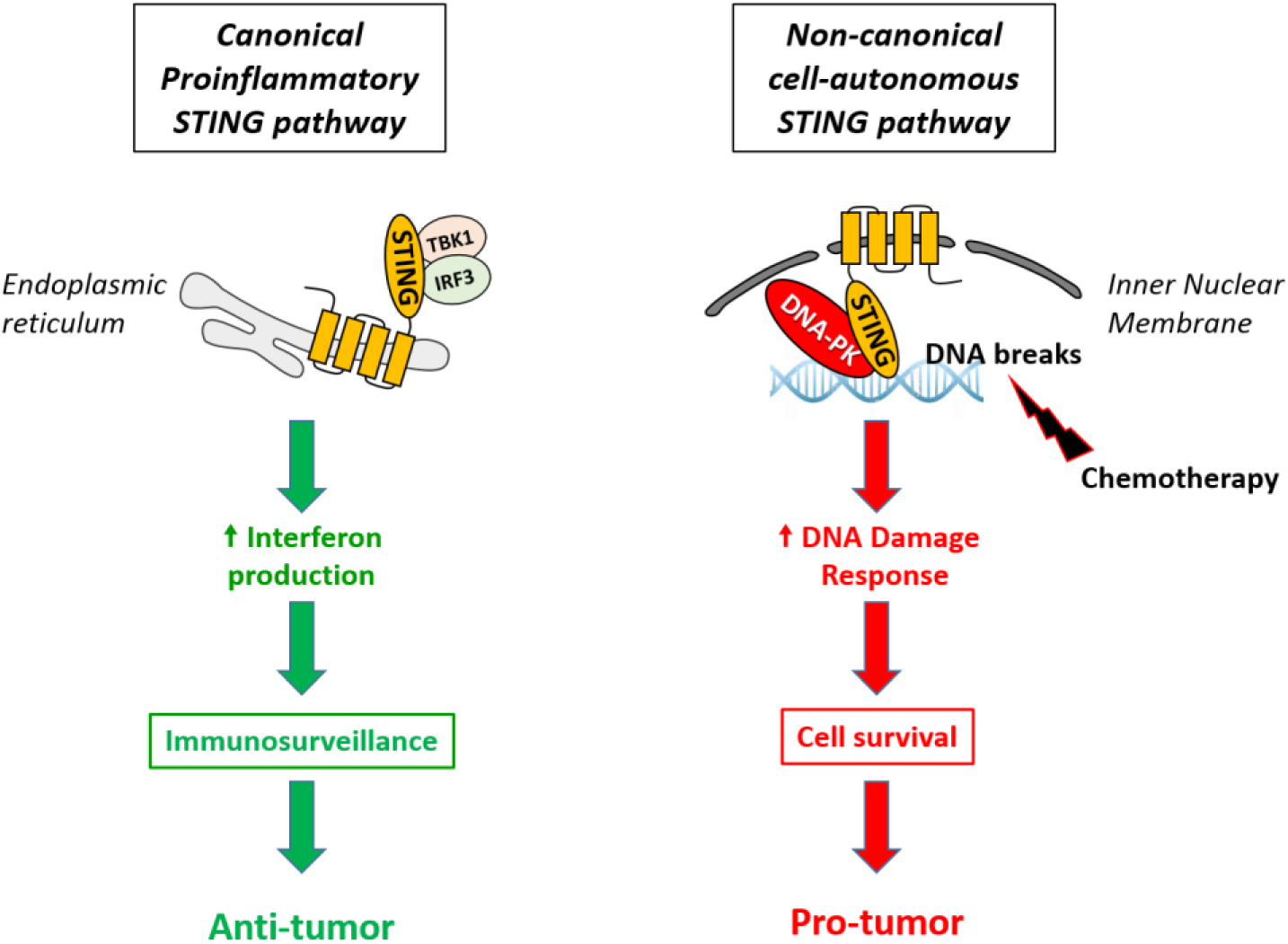

## Introduction

Stimulator of Interferon Genes (STING) has a well-established adaptor function during the innate immune response to cytosolic DNA (Ishikawa, Ma, and Barber 2009). This transmembrane protein is mainly described as an endoplasmic reticulum (ER)-resident protein (Ishikawa and Barber 2008). It is composed of four N-terminal transmembrane domains and a cytosolic C-terminal tail that contains the cyclic dinucleotide (CDN)-binding domain and domains of interaction with downstream effectors (Liu et al. 2015). Upon infection, CDNs are directly secreted by pathogens or generated by the cyclic GMP-AMP synthase (cGAS) in response to cytosolic pathogen-derived DNA (Ablasser et al. 2013). CDNs bind to, and activate, the adaptor protein STING that recruits TANK-binding kinase 1 (TBK1) and the transcription factor IRF-3 (interferon [IFN] regulatory factor-3). TBK1 phosphorylates IRF-3, phospho-IRF-3 forms homodimers that translocate to the nucleus to induce the expression of inflammatory cytokines (Liu et al. 2015).

Genome integrity is constantly threatened by endogenous and environmental genotoxic stresses. DNA damage can arise through the action of reactive oxygen species produced during oxidative metabolism. DNA lesions can also result from exposure to various chemicals and radiation. When DNA integrity is challenged, the cell triggers a complex signaling network called the DNA damage response (DDR), that detects the lesion and organizes its repair (for a review, see Ref Zhou and Elledge 2000). Cells with altered DDR are more prone to develop a variety of diseases, including cancers (Jackson and Bartek 2009). Accordingly, genomic instability is a hallmark of cancer cells in which genomic rearrangements (translocations, deletions, and duplications) are extremely frequent. Many cancer therapies (e.g. radiotherapy, chemotherapy) rely on the induction of DNA damage to drive the killing of rapidly cycling tumor cells (Bouwman and Jonkers 2012, Hosoya and Miyagawa 2014). Alkylating agents, such as nitrogen mustard (e.g. cyclophosphamide) or platinum compounds (e.g. cisplatin), are among the most widely used anti-cancer drugs. They form DNA cross-links that block replication forks, impede cell division, ultimately leading to multiple DNA breaks and cell death (Kondo et al. 2010). As efficient DNA repair pathways can enable tumor cells to survive treatment-induced DNA damage, the identification of new actors contributing to the DDR may help design alternative therapeutic approaches.

We recently linked STING-mediated inflammation to genotoxic stress in the context of breast cancer (Gaston et al. 2016). As cyclophosphamide is commonly used for breast cancer therapy, we treated MCF7 breast cancer cells with mafosfamide, a cyclophosphamide analog suitable for *in vitro* studies as it does not require hepatic activation to generate its active metabolite (4-hydroxy-cyclophoshpamide) (Mazur et al. 2012). Such a genotoxic treatment led to the accumulation of DNA in the cytoplasm of MCF7 cells and triggered the expression of IFNs and of several IFN-stimulated genes via the canonical STING/TBK1/IRF-3 pathway. Similar observations were reported by others using other DNA-damaging agents (e.g. etoposide, cisplatin, cytarabine, irradiation) (Erdal et al. 2017, Ahn et al. 2014, Parkes et al. 2017, Mackenzie et al. 2017, Lan et al. 2014) or even in the context of intrinsic genetic instability characteristic of cancer cells (e.g. BRCA1-mutant breast cancer cells, U2OS osteosarcoma cells) (Parkes et al. 2017, Mackenzie et al. 2017). While STING-mediated inflammation is broadly considered to promote anticancer immune responses (Deng et al. 2014, Vanpouille-Box et al. 2017, Wang et al. 2017, Harding et al. 2017), we showed that abrogation of this inflammatory response *in vitro*, i.e. in absence of a functional immune system, potentiated treatment-induced cell death and delayed cell colony regrowth, suggesting a cell-autonomous contribution of the STING/IFN pathway to the resistance of cancer cells to treatment (Gaston et al. 2016).

Our study suggested that STING may partly reside in the nucleus of MCF7 cells (Gaston et al. 2016). Indeed, biochemical cell fractionation showed that STING was intrinsically present in the nuclear fraction, irrespective of mafosfamide treatment. While STING has been mainly studied as an ER-resident protein, our observations are reminiscent of its original identification as a nuclear envelope transmembrane (NET) protein in liver (hence its original name NET23) (Schirmer et al. 2003). Interestingly, the cytosolic DNA sensor cGAS has been recently shown to translocate to the nucleus upon DNA damage where it contributes to the DDR by suppressing homologous recombination (HR) (Liu et al. 2018). Conversely, some nuclear proteins involved in the DDR (e.g. DNA-PKcs, Ku70 or MRE11) have been shown to act as cytosolic DNA sensor and to activate inflammatory responses (Ferguson et al. 2012, Kondo et al. 2013, Sui et al. 2017, Burleigh et al. 2020). The dual subcellular localization and function of these acknowledged DNA sensors/DDR mediators called for investigating further the nuclear localization of STING and its potential involvement in the DDR.

We here demonstrate that STING partly localizes at the inner nuclear membrane (INM) in breast cancer cells and in patients’ tumors, and promotes the DDR in a CDN/TBK1/IRF3-independent manner. As opposed to its well documented role in stimulating antitumor immunity, our data support that STING intrinsically promotes breast cancer cell survival both in steady-state and upon genotoxic stress via a non-canonical, cell-autonomous mechanism.

## Results

### STING partly resides in the nucleus of breast cancer cells

Our preliminary observations involving MCF7 breast cancer cell line suggested that a part of the cellular STING pool resides in the nuclear fraction (Gaston et al. 2016). To challenge this finding in other models, we investigated a series of so-called HBCx (Human Breast Cancer xenograft) cells generated in-house from breast cancer Patient Derived Xenograft (PDX). STING was detected in all of them (Fig. 1a) at much higher level than in MCF7 cells (Fig 1b). Two day-mafosfamide treatment did not significantly alter STING expression (Fig 1b). In agreement with our former observations (Gaston et al. 2016), STING was detected in the nuclear fraction of MCF7 cells (Fig 1c, left). In high STING-expressing HBCx-3 cells, massive amounts of STING were detected in the nuclear fraction (Fig. 1c, middle). A lower quantity was also detected in the nucleus of HBCx-39 cells that exhibit moderate STING expression (Fig. 1c, right). Since the three breast cancer cell models exhibited similar fractionation profiles (Fig 1c) despite very different expression level of STING (Fig 1b), MCF7 cells were privileged for subsequent studies as they can be more easily manipulated than HBCx models (e.g. more efficient transfection efficiency). To avoid specificity issues encountered in immunocytofluorescence experiments using commercial anti-STING antibodies (Gaston et al. 2019), we generated Flag and/or hemagglutinin (HA)-tagged STING constructs. The latter displayed similar fractionation profile as endogenous STING when ectopically-expressed in MCF7 cells (Fig. 1d) and in STING-deficient HEK293 cells (Fig 1e,f). This indicates that the presence, the nature (HA, Flag) and the position (N- or C-terminal) of tags do not alter the biochemical subcellular localization of STING.

**Fig 1.**
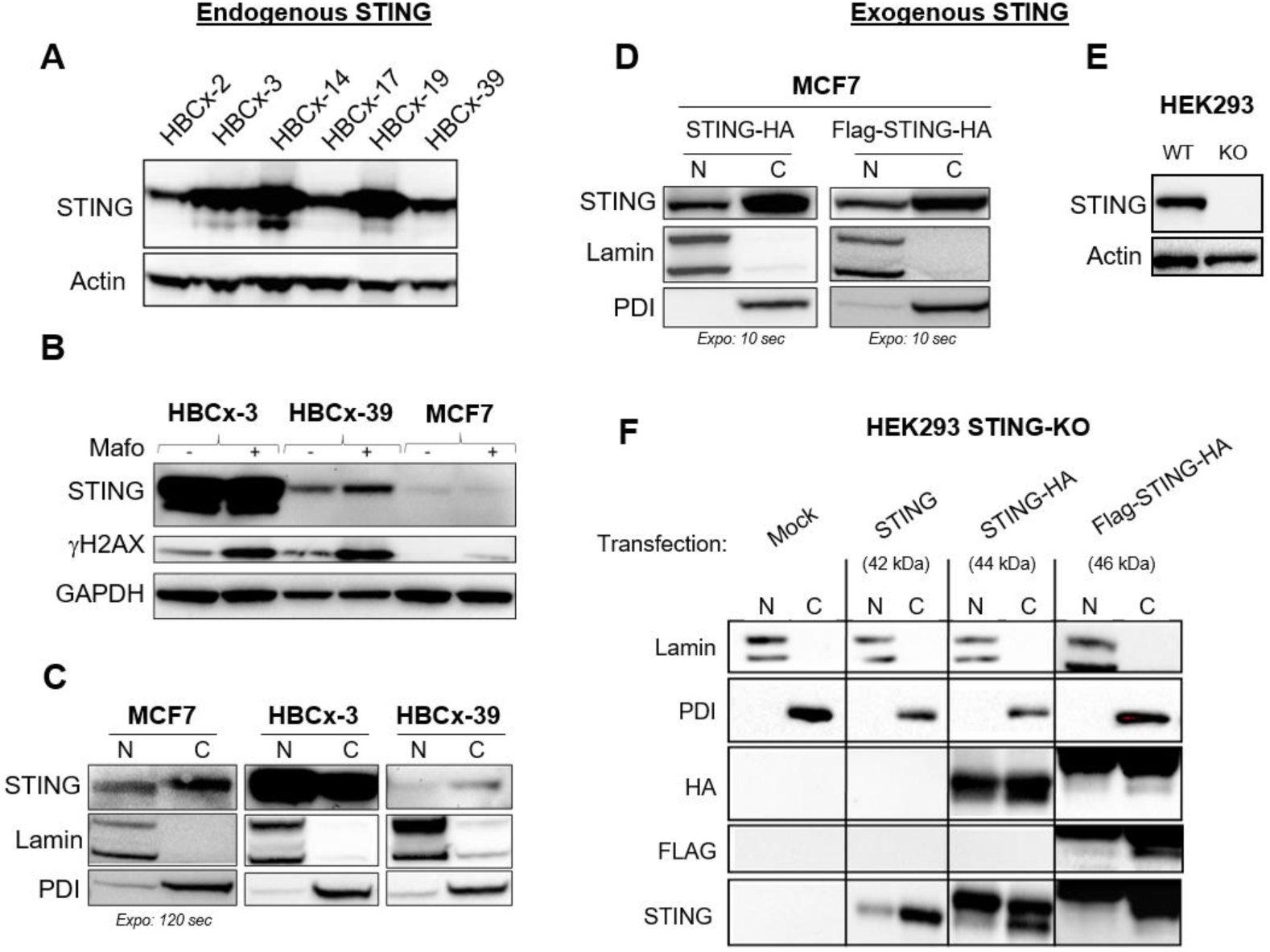
Nuclear localization of STING in various cell lines. **a** Immunoblot of endogenous STING in various PDX-derived breast cancer cell lines (named HBCx). **b** Immunoblot of endogenous STING and γH2AX in HBCx-3 and HBCx-39 *versus* MCF7 cells 48h after treatment with (+) or without (-) mafosfamide (10µM). **c**,**d** Immunoblot of endogenous (**c**) or ectopically expressed (**d**) STING in the cytoplasmic (C) and nuclear (N) cell fractions of breast cancer cells. Lamin A/C is used as a nuclear marker and the ER-resident Protein Disulfide Isomerase (PDI) as the cytoplasmic marker. In MCF7 cells, the time of anti-STING blot exposure was adjusted to the expression level of STING, as indicated. **e** Immunoblot of endogenous STING in parental *versus* STING-KO HEK293 cells. **f** Immunoblot of lamin A/C, PDI and different tagged STING constructs in the cytoplasmic (C) and nuclear (N) fractions of transiently transfected STING-KO HEK293 cells.

Taken together, these data show that part of the STING pool resides in the nuclear fraction of various cell types. When STING is ectopically expressed, this detection in the nuclear fraction is maintained irrespective of tags.

### STING co-localizes with the lamina in breast cancer cells

We further characterized STING subcellular localization using immunofluorescence staining. As shown in Fig. 2a, STING was uniformly spread within the cytoplasm of MCF7 cells, and, as expected, partially co-localized with ER (calnexin) and Golgi (GM130) markers (Saitoh et al. 2009). Strikingly, STING also co-localized with lamin B1, a component of the nuclear lamina (Fig 2a, right). The lamina is a fibrillary network underlying the inner nuclear membrane (INM) that serves as anchoring point for INM proteins, chromatin and transcription factors (Dobrzynska et al. 2016). To further investigate this finding, we performed a pre-fixation ribonuclease- and detergent-based cell extraction that preferentially retains cytoskeleton, nuclear matrix and chromatin, at the expense of soluble/loose structures (Britton, Coates, and Jackson 2013, Sawasdichai et al. 2010). The perinuclear ER and nuclear lamina, but not the Golgi apparatus, were preferentially retained after this treatment (Fig. S1a). STING strongly co-localized with lamin B1 (Fig. 1b) and lamin A/C (Fig. S1b) at the nuclear rim (white arrowheads) and at small intranuclear structures (white arrows) presumably corresponding to nuclear membrane invaginations.

**Fig 2.**
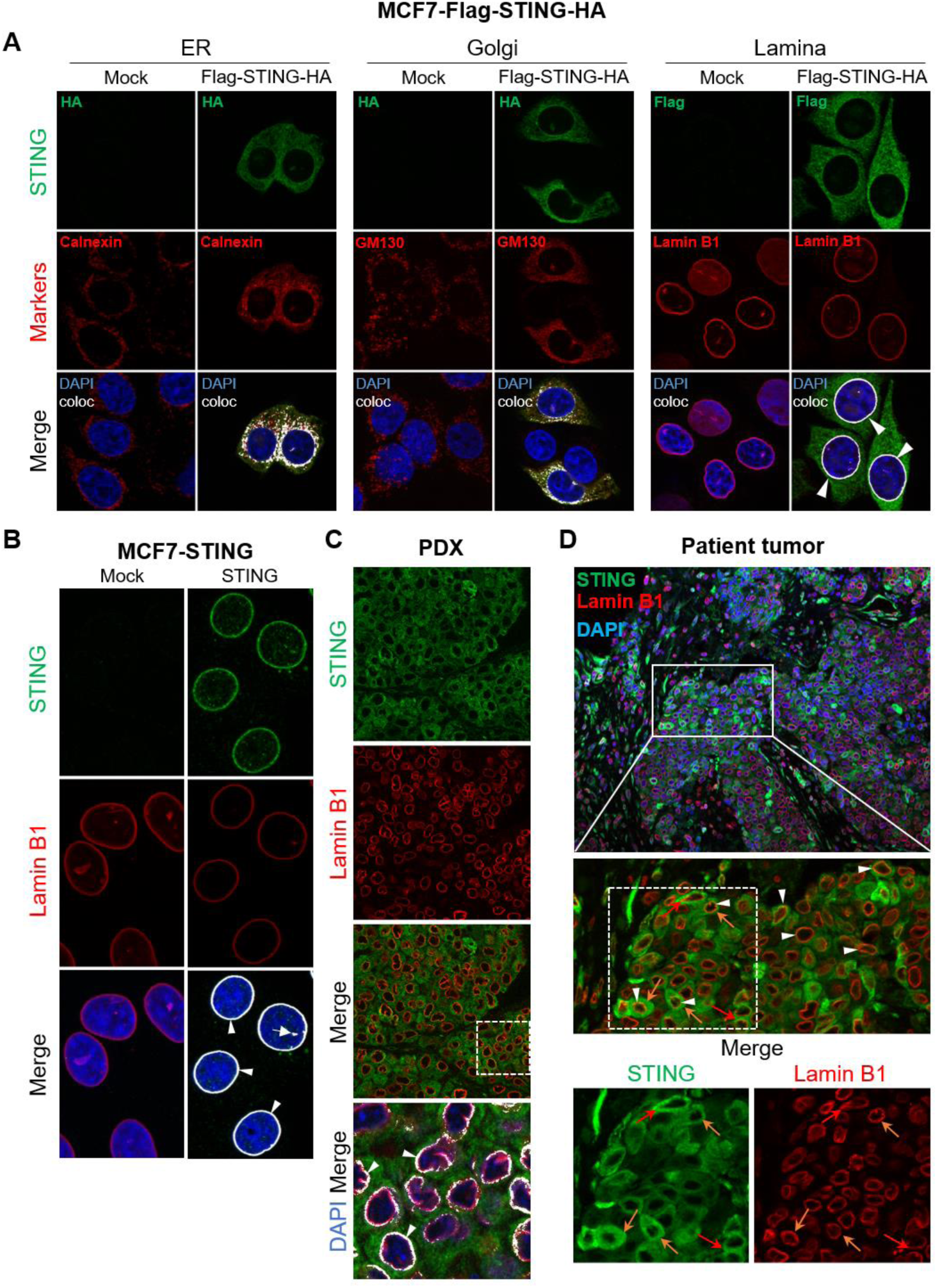
Nuclear STING co-localizes with the lamina in breast cancer cells and tumors. **a** Immunofluorescence of Flag-STING-HA transiently expressed in MCF7 cells (*versus* empty vector, mock), using anti-HA or anti-Flag antibodies (according to the species of the antibody directed against subcellular markers) as indicated (upper panels). Middle panels show immunofluorescence of ER (calnexin), Golgi (GM130) and nuclear lamina (lamin B1) markers. Lower panels display merged images. **b** Immunofluorescence experiment performed after pre-extraction of MCF7 cells transfected with untagged STING (*versus* mock vector), using anti-STING and anti-lamin B1 antibodies. Lower panels display merged images. In **a** and **b**, white arrowheads point to co-localization of STING at the lamina rim, and white arrows to intra-nuclear staining. Nuclei were stained with DAPI. **c** Immunofluorescence of endogenous STING and lamin B1 in the high STING-expressing HBCx-19 PDX. Lower panels display merged images with or without DAPI and image treatment by Image J software (shown as an inset at higher magnification) to emphasize co-localization (appearing in white). **d** The upper image shows one representative area of a patient tumor immunostained for STING and lamin B1 (nuclei stained with DAPI). The three bottom panels show higher magnification of the squared area for STING, lamin B1 and merged staining, as indicated. White arrowheads: examples of STING/lamin B1 co-localization; orange arrows: cells in mitosis. See also Fig S1 and S2.

To address the clinical relevance of our findings, we investigated the nuclear localization of STING in malignant breast tumors. First, we analyzed 4 breast cancer PDXs (estrogen receptor-positive [ER+] and triple negative [TN] subtypes) exhibiting different tissue levels of STING mRNA (high, medium, low; data not shown). Accordingly, different levels of STING protein were detected in tumor cells by immunofluorescence, assessing immunostaining specificity (Fig. S2a). In high STING-expressing samples, a clear co-localization of STING with lamin B1 was observed at the nuclear rim (Fig 2c and Fig. S2b). Second, we analyzed samples of 6 TN breast cancers resistant to neoadjuvant treatment containing cyclophosphamide (representative examples are shown in Fig. 2d and Fig. S2c). STING was detected in all samples analyzed, and in each sample the staining for STING, of various intensities, was present in the tumor cells. The STING/lamin B1 co-localization was observed at the nuclear rim of tumor cells (Fig 2d and Fig. S2c, white arrowheads). Some of those cells were in mitosis (Fig 2b, orange arrows).

Together, these data demonstrate that in various cell lines, PDXs and clinical tumor samples, a fraction of the STING pool intrinsically co-localizes with the lamina in the nucleus of breast cancer cells.

### Identification of STING at the INM by electronic microscopy

Considering the transmembrane nature of STING, its co-localization with the lamina, and the resistance of INM proteins to pre-extraction (Malik et al. 2010), we hypothesized that nuclear STING localizes at the INM. To explore this hypothesis further, we monitored STING localization by immunogold labeling in immunoelectron microscopy (EM). STING protein was stained using anti-flag antibody and nanogold secondary antibody in MCF7 cells expressing Flag-STING-HA construct (Fig. 3a). Immunogold staining specificity was confirmed using non-transfected cells as a control (Fig. S3a). As shown in Fig. 3b-d, black dots corresponding to anti-Flag-bound gold particles were observed at cytoplasmic vesicle structures (white arrowheads) and, in agreement with STING/calnexin co-localization, at perinuclear ER membranes (grey arrowheads). Consistent with our immunofluorescence results, STING was detected at the periphery of the nucleus, mainly at proximity of the INM at a distance compatible with immunogold staining of a transmembrane protein (black arrowheads, Fig. 3c-f). Furthermore, STING was frequently observed at both sides of nuclear membrane invaginations (Fig. 3f) and sometimes appeared as gold dot doublets (Fig. 3e,f) that likely represent distinct quaternary structures of STING as recently characterized by cryo-EM (Shang et al. 2019).

**Fig 3.**
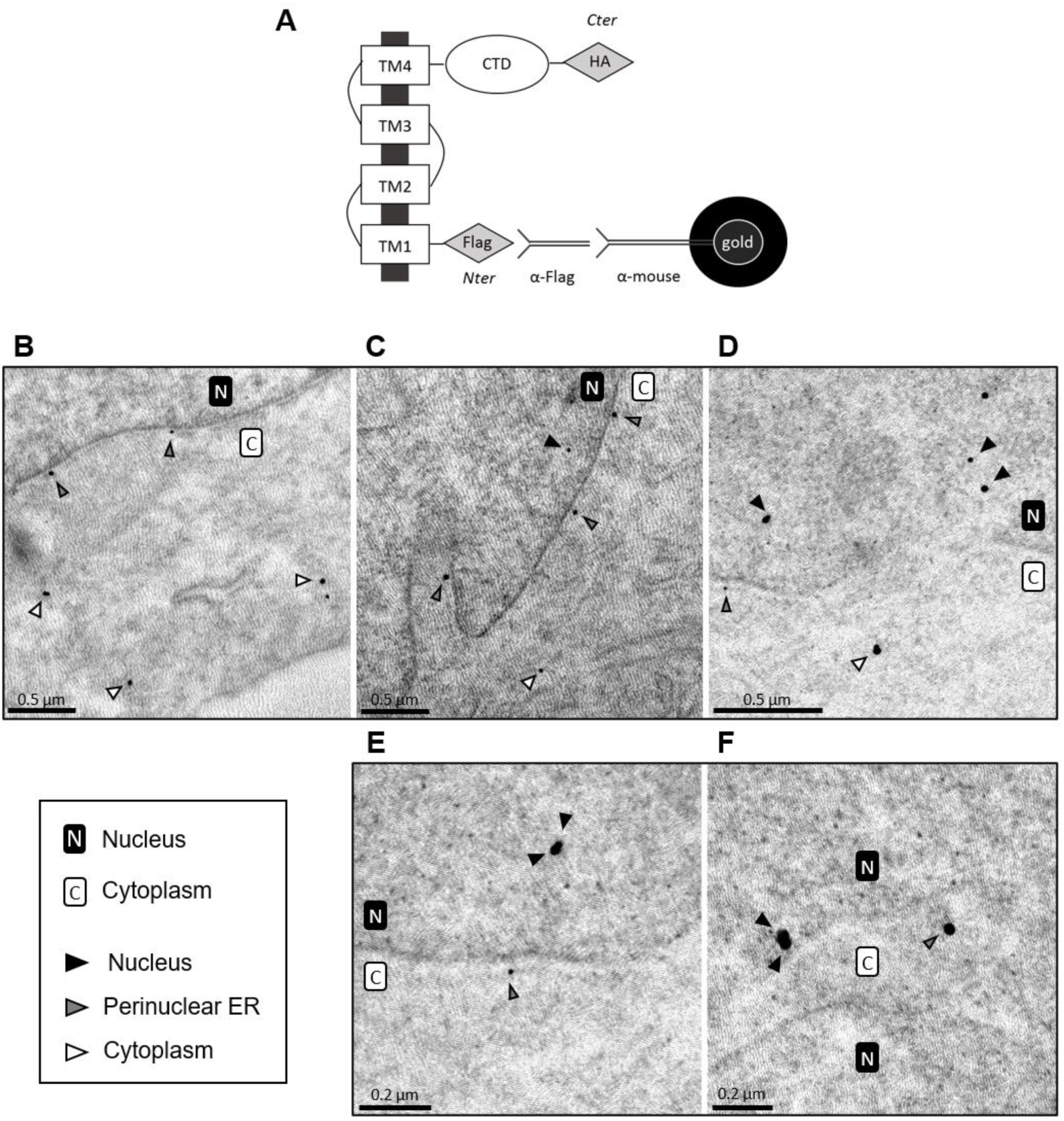
STING localizes at the INM. **a** Schematic representation of the anti-Flag immunogold staining procedure of MCF7 cells expressing the Flag-STING-HA construct. **b-f** Five representative immunoelectron microscopy images illustrating typical features of STING subcellular localization identified using symbols displayed in the bottom left box. Negative control involved parental cells (Fig. S3a). Size bars are indicated in each panel. See also Fig S3.

Cells analyzed 48h after mafosfamide genotoxic treatment showed several signs of stress including nuclei with irregular shape and large invaginations, picnotic nuclei, dilated ER with dramatically enlarged lumen and large vesicles filled with cell debris (Fig. S3b, bottom panels). As previously reported (Gonugunta et al. 2017), many cytoplasmic vesicles were positive for STING (Fig. S3b, upper left panel). The various locations of STING reported above for naïve cells were also observed in mafosfamide-treated cells (Fig. S3b, upper panels).

Taken together, these data demonstrate that in breast cancer cells, the nuclear STING pool mainly resides at the nucleus periphery and at the INM.

### STING promotes the DDR

There is accumulating evidence for dual function of proteins involved in the response to cytosolic DNA and the DDR. The finding that STING partly localizes in the nucleus of breast cancer cells triggered us to inquire about the potential involvement of STING in the DDR. To investigate this hypothesis, we manipulated the levels of STING expression and assessed the consequences on DDR efficiency. The formation of 53BP1 (p53 Binding Protein 1) and γH2AX (phosphorylated H2AX histone) foci at DNA damage sites is a hallmark of DDR initiation (Schultz et al. 2000). STING loss-of-function (LOF) using targeted (*versus* non-targeted [shNT]) shRNA (Fig. S4a) significantly reduced the formation of 53BP1 foci in naïve MCF7 cells, and this effect was significantly rescued by STING overexpression (Fig. 4a,b). Similar results were observed when we compared parental *versus* STING-KO HEK293 cells (Fig. 4c and Fig. S4b). We then quantified DNA damage (single stranded (SSB) and double stranded (DSB) DNA breaks) using the denaturing comet assay (Fig. 4d). Compared to mafosfamide-treated shNT-MCF7 cells, STING LOF (using two different shRNAs) doubled the tail moment and this effect was significantly rescued by STING transient expression (Fig. 4d,e and Fig. S4c). Similar results were obtained in the absence of genotoxic stress in HEK293 and in MCF7 cells (Fig. 4f). Of interest, in naïve MCF7 cells, STING partly co-localized with 53BP1 foci, mainly at the periphery of the nucleus, and this effect was amplified under mafosfamide treatment (Fig. 4g). Partial co-localization was also observed with γH2AX foci (Fig. 4h).

**Fig 4.**
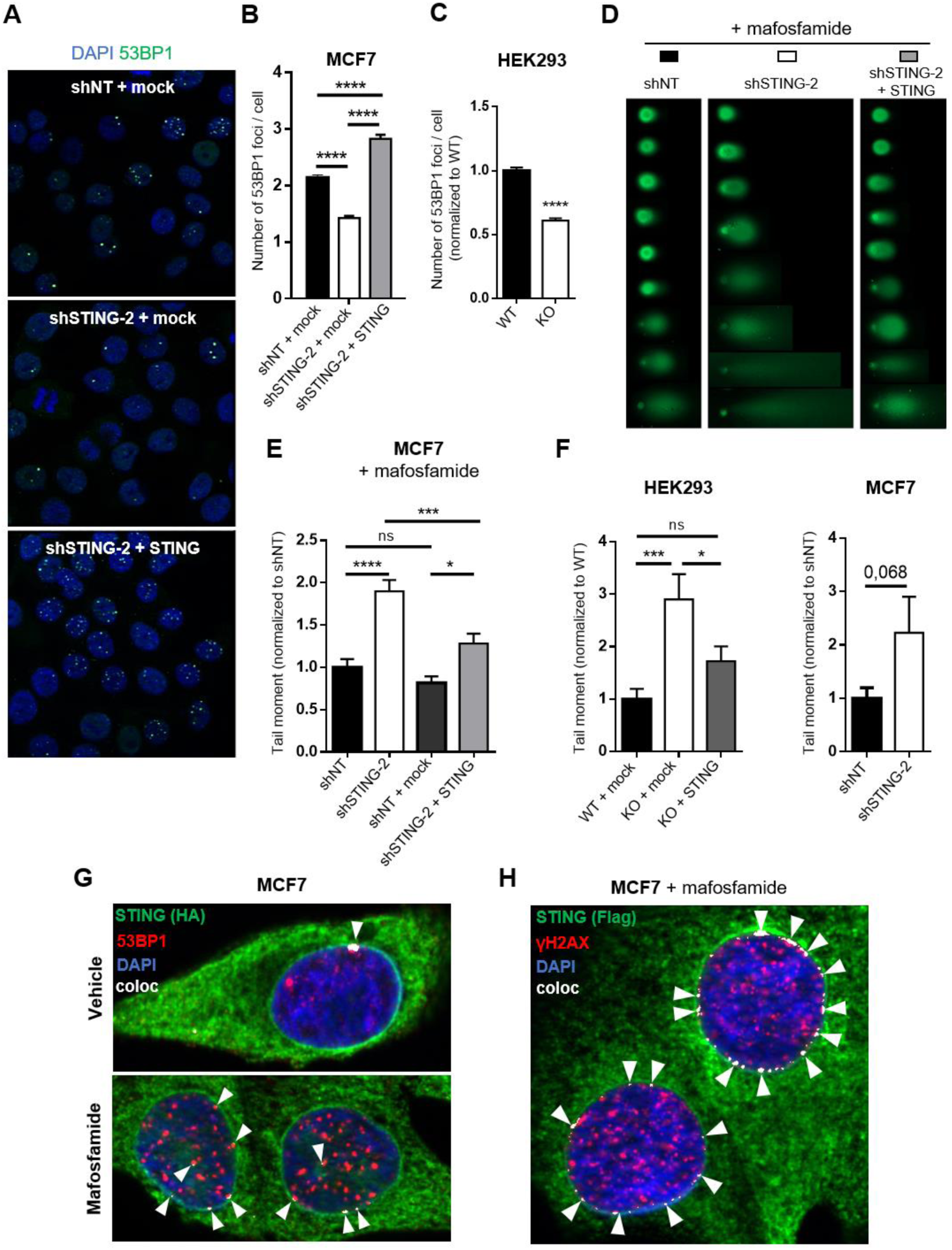
STING contributes to the DDR. **a** Immunofluorescence of endogenous 53BP1 foci in non–treated MCF7 cells transduced with shNT or shSTING-2 and rescued or not for STING expression, as indicated. Nuclei were stained with DAPI. Quantification in **b**: mean ± s.e.m. of the number of 53BP1 foci per cell of n= 2,304 (shNT + empty vector), n= 2,482 (shSTING + empty vector) and n= 2,276 (shSTING + STING vector) cells from n=3 independent experiments (one-way ANOVA and post-hoc Tukey’s multiple comparisons test). **c** Quantification of endogenous 53BP1 foci in treatment-naïve parental (WT) versus STING-KO HEK293 cells. Mean ± s.e.m of n=1,489 (WT) and n=1,419 (KO) cells from 3 independent experiments (Student’s t-test). **d-f** Comet assays. Representative images of comet assays (**d**) and quantification of the tail moment with (**e**) or without (**f**) mafosfamide treatment of MCF7 cells transduced with shSTING-2 versus shNT, and HEK293 cells STING-KO versus WT, rescued (STING) or not (mock) for STING expression, as indicated. In **e**: mean ± s.e.m. of tail moment of n= 746 (shNT), n= 976 (shSTING-2), n= 776 (shNT + mock plasmid) and n= 605 (shSTING-2 + STING vector) cells from n=3 independent experiments (one-way ANOVA and post-hoc Tukey’s multiple comparison test). In **f:** mean ± s.e.m of tail moment of n=91 (HEK WT + mock), n=89 (HEK-KO + mock), n=117 (HEK-KO + STING), n=132 (MCF7-shNT) and n=118 (MCF7-shSTING-2) cells (HEK cells: one-way ANOVA and post-hoc Tukey’s multiple comparison test, MCF7 cells: Student’s t-test). **g**,**h** Immunofluorescence analysis of STING, 53BP1 and γH2AX foci. Colocalization (arrowheads) of STING with 53BP1 (**g**) and γH2AX (**h**) in MCF7-Flag-STING-HA cells treated for 48h with vehicle or mafosfamide, as indicated. See also Fig S4.

Taken together, these data demonstrate that STING promotes the DDR in basal and in genotoxic-induced stress conditions.

### STING promotes intrinsic breast cancer cell survival and resistance to genotoxic stress

Several recent studies reported that STING-mediated cytokine production promoted anticancer immune responses (Deng et al. 2014, Vanpouille-Box et al. 2017, Wang et al. 2017, Harding et al. 2017). However, the involvement of STING in the DDR uncovered in this study suggests a potential cell-autonomous contribution of STING to cancer cell survival.

To address this issue, we investigated whether manipulation of STING expression impacted cancer cell survival *in vitro*, i.e. in absence of a functional immune system. Clonogenic survival assays were used to address the ability of cancer cell to survive and resume proliferation after genotoxic treatment. In MCF7 and BT20 breast cancer cell lines, STING depletion significantly reduced the number of mafosfamide-resistant clones observed 40 days post-treatment (Fig 5a,b). Conversely, transient STING overexpression at the time of drug addition markedly enhanced resistance to treatment as reflected by the higher number of clones at day 20 (Fig 5c). To broaden these findings, we tested various experimental conditions using standard cell viability assays as readouts. First, STING expression was transiently silenced in various breast cancer cell models using siRNA (Fig 5d), and cell viability (*versus* siNT-treated cells) was measured 10 days later. STING inhibition drastically reduced cell viability in various ER+ and TN breast cancer models including MCF7 (Fig 5e), BT20, HCC1937 and four HBCx cell lines (Fig. 5f). In HBCx cells, cell survival was inversely proportional to siRNA efficiency (due to poor transfection efficacy of some models). Second, we performed viability assays under genotoxic conditions. These experiments were performed using stable STING-deficient systems in order to circumvent any bias due to the intrinsic toxicity of transient transfection procedures. In addition to mafosfamide, which generates DNA breaks by cross-linking nucleotides, we also investigated the response to Etoposide, another class of genotoxic agent that stabilizes transient DSB via inhibition of the topoisomerase II. Although Etoposide is irrelevant to breast cancer therapy, it is frequently used in DNA damage studies. In both MCF7 and HEK293 cells and irrespective of the genotoxic agent, STING deficiency significantly increased drug-sensitivity. Although of moderate amplitude, this effect was highly robust (Fig 5g-j).

**Fig 5.**
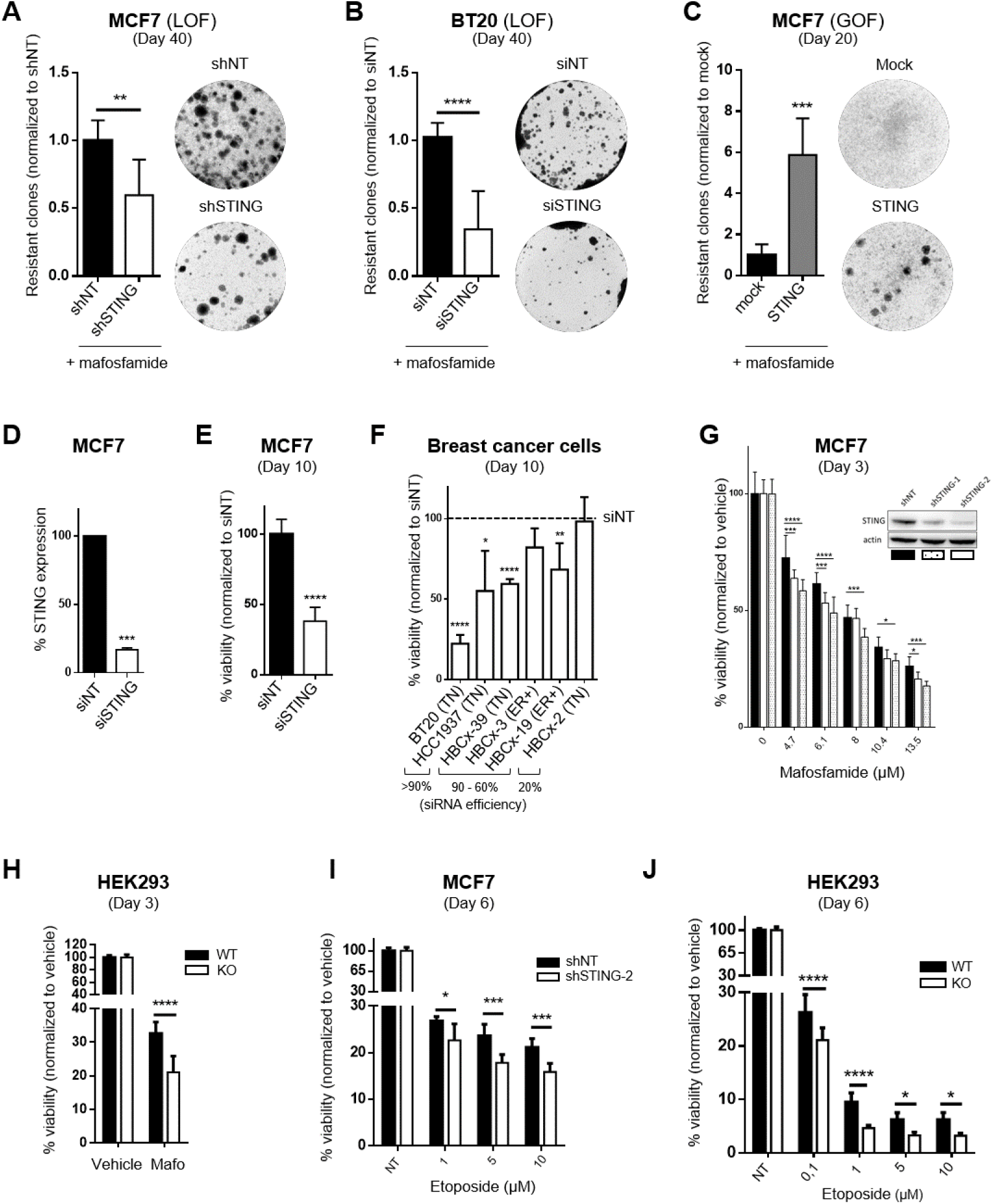
STING promotes intrinsic breast cancer cell survival and resistance to genotoxic stress. **a-c** GOF and LOF colony assays of MCF7 (**a**,**c**) and BT20 (**b**) cells showing the regrowth of cells 40 days (**a**,**b**) or 20 days (**c**) after exposure to mafosfamide (10 µM). MCF7 cells were stably transduced with shNT or shSTING prior to treatment (**a**) and rescued or not (mock vector) (**c**) by the transfection of a STING-encoding plasmid the day of mafosfamide treatment. In BT20 cells (**b**), STING was silenced by siRNA 3 days before mafosfamide exposure. Representative wells are shown. Mean ± s.d. of n=9 from 3 independent experiments, Student’s t-test. **d-f** Effect of transient STING silencing in naïve breast cancer cells. **d** RT-qPCR analysis showing the efficacy of a siRNA targeting *STING* (siSTING) *versus* a non-targeted siRNA (siNT) (Student’s t-test, n=2 independent experiments). **e**,**f** Viability of MCF7 cells (**e**) (Mean ± s.d. of n=9 from 3 independent experiments, Student’s t-test) and of two immortalized and four PDX-derived breast cancer HBCx cells (**f**) (Mean ± s.d of n=6 from 2 independent experiments, Student’s t-test) 10 days after transfection with siNT *versus* siSTING. In **f**, the molecular subtype (ER, TN) and the efficiency of siRNA to inhibit *STING* expression is indicated. **g-j** Effect of stable STING silencing on MCF7 and HEK293 cells sensitivity to genotoxic treatment. **g** Cell viability of MCF7 cells stably transduced with either shNT or shRNAs targeting *STING* was measured 3 days after treatment with various doses of mafosfamide (mean ± s.d. of n=9 from 3 independent experiments, two-way ANOVA and post hoc Dunnett’s multiple comparison test). **h** Cell viability of parental HEK293 *versus* STING-KO HEK293 cells 3 days after exposure to 5 µM mafosfamide (mean ± s.d. of n=10 from 2 independent experiments, two-way ANOVA and post hoc Sidak’s multiple comparison). **i**,**j** Same as in **g** and **h** but 6 days after exposure to various doses of etoposide, as indicated (mean ± s.d. of n=12 from 2 independent experiments, two way ANOVA and post hoc Sidak’s multiple comparison).

Together, these data demonstrate that STING promotes breast cancer cell survival and contributes to resistance to genotoxic stress in a cell-autonomous manner.

### STING-mediated promotion of DDR and cell survival is independent of its canonical pro-inflammatory pathway

The canonical STING/TBK1/IRF-3/IFN pathway has been shown by us and others to be activated in various preclinical breast cancer models in response to genotoxic stress (Li and Chen 2018, Legrier et al. 2016, Gaston et al. 2016, Erdal et al. 2017, Parkes et al. 2017). This raised the question whether this inflammatory pathway could interfere with the novel function of STING discovered in the present work. The fact that STING contributes to the DDR in the absence of genotoxic stress, i.e. in culture conditions lacking IFN induction (Gaston et al. 2016), was against this hypothesis. Nonetheless, we aimed to address this issue experimentally. First, exogenous activation of the STING/TBK1/IFN pathway using the CDN cGAMP (a typical STING agonist; see Ref. (Ablasser et al. 2013)) expectedly triggered IFN signaling as reflected by the upregulation of typical IFN-stimulated genes (Fig 6a), but had no impact on the formation of 53BP1 foci (Fig. 6b). Second, in contrast to STING LOF (Fig. 4b), TBK1 silencing (Fig. 6c) did not impair the formation of 53BP1 foci, that was even slightly increased (Fig. 6d). Third, a naturally occurring C-terminally truncated STING isoform has been shown to act as a dominant-negative (DN) of full-length STING on IFN induction due to its inability to interact with TBK1/IRF3 complex (Chen et al. 2014). Cell fractionation analyses showed that STING-DN displayed similar subcellular distribution compared to full-length STING, although the former tended to be predominant in the nucleus fractions (Fig. 6e). Expression of STING-DN in a shRNA-mediated-STING-deficient background was sufficient to protect MCF7 cells from DNA damage accumulation as revealed by the comet assay (Fig. 6f). Fourth, as opposed to STING silencing, IFN receptor (IFNAR1) silencing (Fig. 6g) had no effect on MCF7 cell survival in steady-state as well as 10 days after mafosfamide treatment (Fig. 6h), i.e. when IFN production has been triggered (Gaston et al. 2016). Taken together, these data strongly suggest that STING promotes the DDR and cancer cell survival in a cell-autonomous and CDN/TBK1/IFN-independent manner.

**Fig 6.**
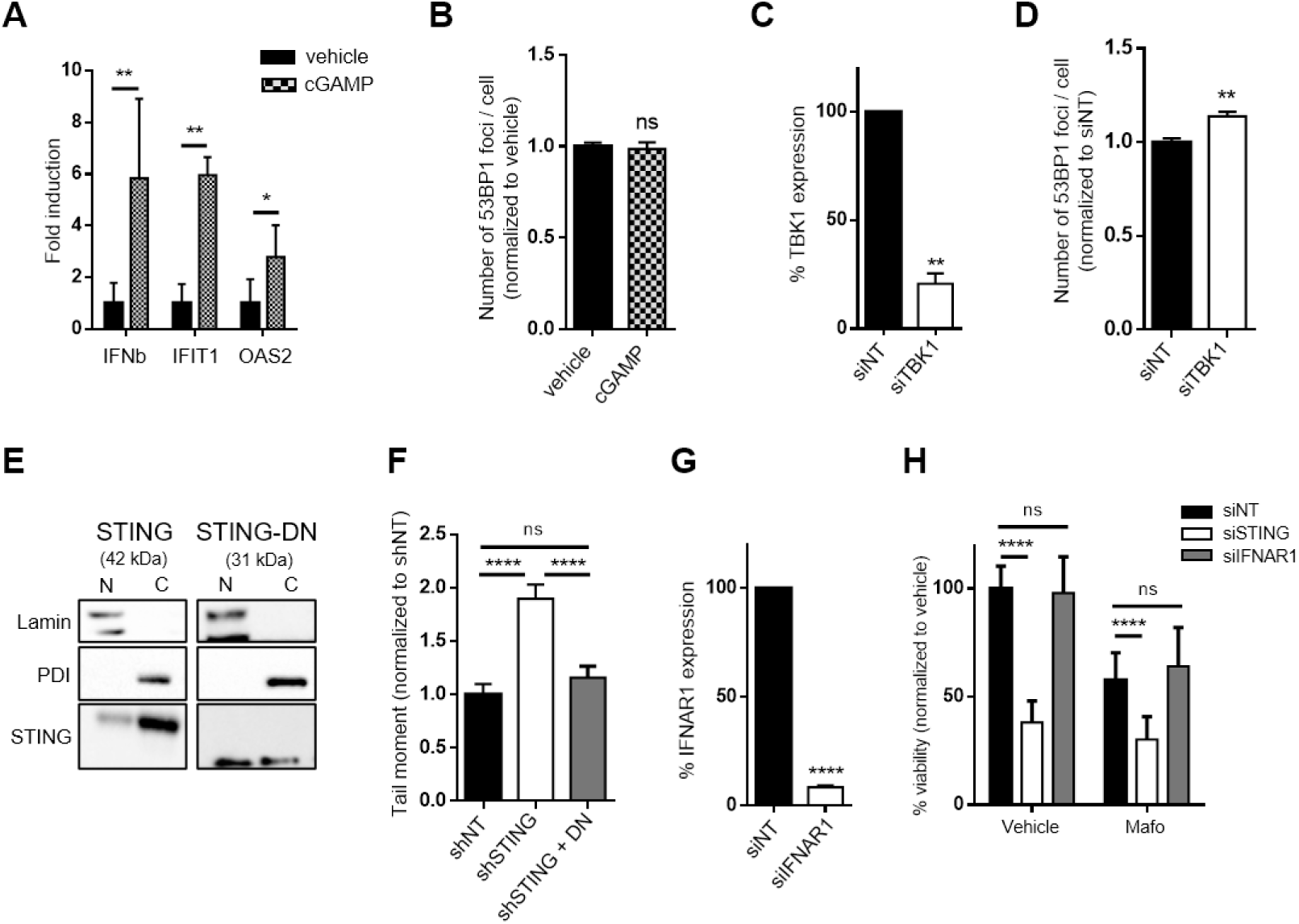
The effects of STING on the DDR and cell survival are independent of the canonical inflammatory pathway. **a** RT-qPCR analysis of *IFNβ, IFIT1 and OAS2* expression in MCF7 cells 6h after exposure to the CDN cGAMP. Mean ± s.d. of n=3 independent experiments, Student’s t-test. **b** Effect of cGAMP treatment on the formation of 53BP1 foci in MCF7 cells as determined by immunofluorescence. Mean ± s.e.m of the number of foci per cell in n=3,134 (vehicle) and n=3,204 (agonist) cells from n=3 independent experiments (Student’s t-test). **c** RT-qPCR analysis of endogenous TBK1 in MCF7 cells transfected with a siRNA targeting TBK1 (siTBK1) *versus* a non-targeted siRNA (siNT) (Student’s t-test, n=2 independent experiments). **d** Effect of TBK1 silencing on the formation of 53BP1 foci in MCF7 cells as determined by immunofluorescence. Mean ± s.e.m of the number of foci per cell in n= 2640 (shNT) and n= 2340 (siTBK1) cells from n=3 independent experiments (Student’s t-test). **e** Immunoblot of STING and STING-DN in cytoplasmic (C) and nuclear (N) fractions prepared from transiently transfected MCF7 cells. **f** Tail moment of MCF7 stably transduced with shNT or shSTING-2 with or without rescue using a STING-DN expression vector. Mean ± s.e.m. of tail moment of n= 746 (shNT), n= 976 (shSTING-2), and n= 871 (shSTING-2 + STING-DN plasmid, STING-DN) cells from n=3 independent experiments (one-way ANOVA and post-hoc Tukey’s multiple comparison test). **g** RT-qPCR analysis of endogenous *IFNAR1* expression in MFC7 cells transfected with a siRNA targeting *IFNAR1* (siIFNAR1) *versus* a non-targeted siRNA (siNT) (Student’s t-test, n=2 independent experiments). **h** Viability of MCF7 cells 10 days after transfection with siNT, siSTING or siIFNAR1 exposed (right) or not (left) to mafosfamide (10 µM) 3 days after transfection. Mean ± s.d of n=9 from 3 independent experiments, two way ANOVA and post hoc Dunnett’s multiple comparison test.

### Determination of nuclear STING interactome using mass spectrometry

To shed light on the potential mechanism by which STING could promote the DDR, we performed an interactomics analysis. STING was immunoprecipitated from nuclear extracts that were treated with Benzonase beforehand to avoid DNA- or RNA-mediated co-precipitation (Fig. S5a), and the proteins eluted were identified and quantified by mass spectrometry. We confirmed that immunoprecipitates were significantly enriched for STING and nuclear proteins (Fig. S5b and Table S1). Interestingly, none of the canonical STING interactors (e.g. TBK1, IRF3, MAVS or STAT6) (Ishikawa and Barber 2008, Chen et al. 2011, Liu et al. 2015) could be identified in nuclear STING interactome (Table S1). Functional analysis of the nuclear proteins specifically immunoprecipitated by STING revealed the enrichment of three functional networks: DNA-repair, mRNA splicing and eukaryotic translation elongation (STRING database, Fig. 7a).

**Fig 7.**
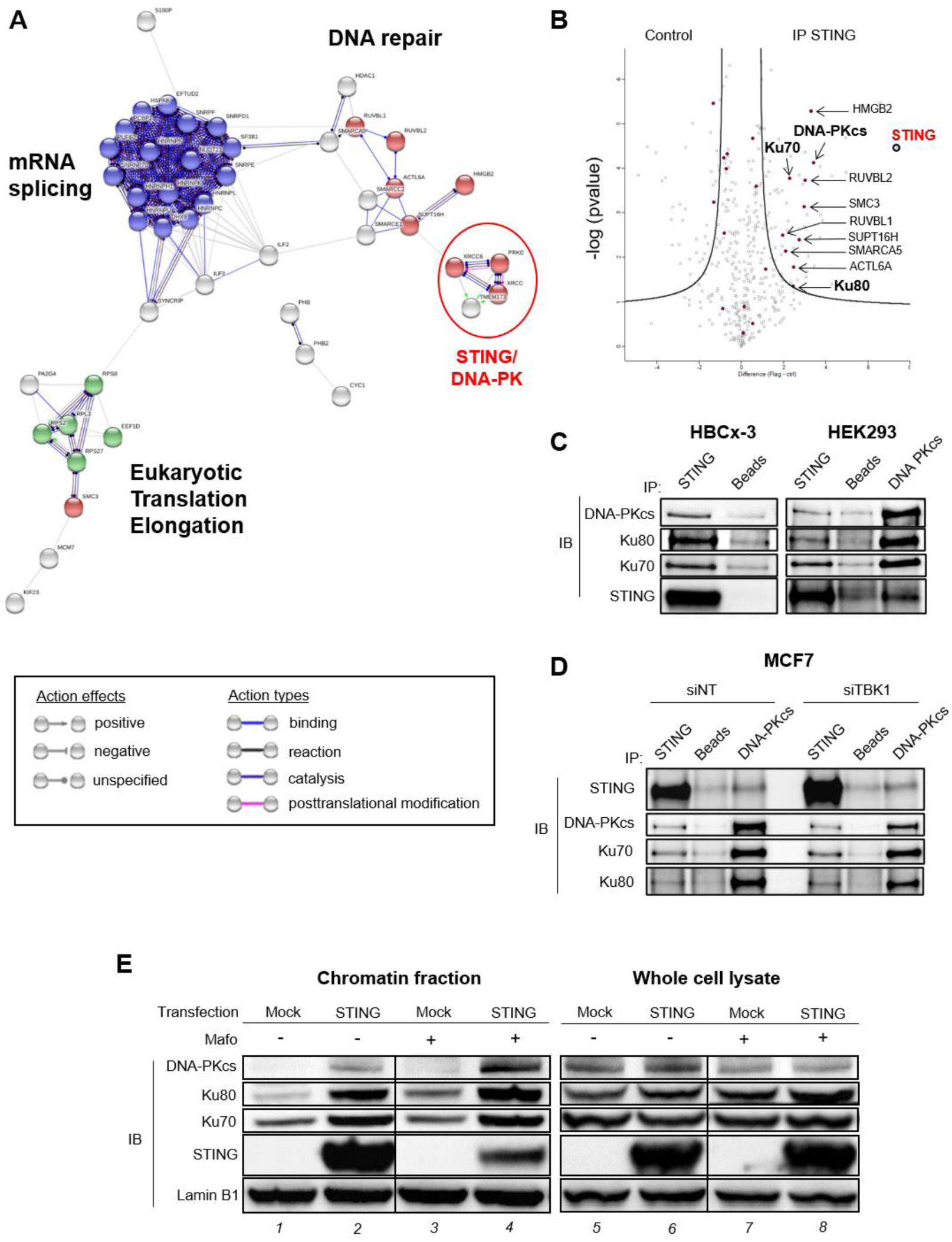
Nuclear STING interactome by mass spectrometry. **a** Functional networks of nuclear STING interactome using STRING database (interaction score high evidence=0,700). Proteins involved in mRNA splicing (FDR=1.22e-18) and Eukaryotic Translation Elongation (FDR=0.00025) pathways, as per Reactome database, are highlighted in blue and green, respectively. Proteins involved in DNA repair according to GO terms (FDR=0.00016) are highlighted in red. **b** Volcano plot of –log(*p*value) *versus* fold change (expressed in log2) of proteins present in anti-STING immunoprecipitates *versus* negative control as detected by mass spectrometry. Proteins previously reported to be involved in DNA repair (Gene Ontology Cell Component database) are colored in magenta. **c** Immunoblots (IB) of proteins constituting the DNA-PK complex (DNA-PKcs, Ku80, Ku70) in immunopecipitates of endogenous STING expressed in HBCx-3 and HEK293 cells. The reverse immunoprecipation was performed in HEK293 cells using anti-DNA-PKcs antibody (right lane). **d** Effect of TBK1 silencing on STING/DNA-PK complex formation in MCF7 cells as assessed by co-immunoprecipitation. In **c-d**, the negative control involved beads only. **e** Immunoblot of proteins constituting the DNA-PK complex (DNA-PKcs, Ku80, Ku70) in the chromatin fractions (lanes 1-4) *versus* whole cell lysates (lanes 5-8) of MCF7 cells overexpressing untagged STING or not (mock) and treated (+) or not (-) with mafosfamide (10µM) for 48h. Lamin B1 is used as a loading control. See also Fig S5.

Given the contribution of STING to the DDR uncovered in this work, we focused on proteins of the DNA repair network. The latter contains many proteins involved in chromatin remodeling complexes that facilitate the efficacy of DNA damage signaling and/or repair, e.g. SMARCA5 (Smeenk et al. 2013), ACTL6A (Lans, Marteijn, and Vermeulen 2012), SUPT16H (Oliveira et al. 2014), RUVBL1/2 (Clarke et al. 2017), SMC3 (Potts, Porteus, and Yu 2006) and HMGB2 (Shin et al. 2013) (Fig 7a, b). However, the most striking observation was the identification of the three core proteins forming the DNA-dependent protein kinase (DNA-PK) complex as part of STING interactome: DNA-PK catalytic subunit (DNA-PKcs), Ku70 (aka XRCC6) and Ku80 (aka XRCC5) (Fig. 7a, b). DNA-PK, together with ATM and ATR, is a master regulator of the DDR (Blackford and Jackson 2017). DNA-PK is mostly known for its involvement in DSB repair through promotion of the Non-Homologous End Joining (NHEJ) repair pathway (reviewed in Ref. Neal and Meek 2011). It is also involved in Homologous Recombination (HR) repair pathway (Neal et al. 2011), checkpoint activation (Liu et al. 2012) and transcription regulation (Pankotai et al. 2012, Calkins, Iglehart, and Lazaro 2013) following DNA damage. This prompted us to investigate further the relevance of DNA-PK regarding nuclear STING.

STING and DNA-PK interaction was confirmed by co-immunoprecipitation experiments involving benzonase-treated extracts from i) tagged (Supplemental Fig. S5c) and untagged (Supplemental Fig. S5d) STING ectopically-expressed in MCF7 cells, ii) endogenous STING in HBCx-3 and HEK293 cells (Fig. 7c), and iii) DNA-PKcs immunoprecipitations showing STING enrichment in addition to Ku70/80 (right lanes of Fig. 7c and Supplemental Fig. S5d). Importantly, TBK1 silencing did not impair STING and DNA-PK interaction, further arguing for the independence of nuclear STING pathway from the canonical cytosolic inflammatory pathway (Fig 7d). Finally, we examined whether STING impacts DNA-PK complex assembly on chromatin using a previously described protocol (Ochi et al. 2015). As shown in Fig. 7e, STING overexpression markedly enhanced the amount of chromatin-bound DNA-PK complex proteins (lane 2 vs 1) without affecting their expression at the cellular level (lane 6 vs 5). In agreement with genotoxic-induced DNA damage, the amount of chromatin-associated DNA-PK complex increased upon mafosfamide treatment and this was further enhanced in the context of STING overexpression (lane 4).

Together, these pioneering observations suggest that STING may cooperate with the DDR regulator DNA-PK, both in basal and in genotoxic-induced stress conditions.

## Discussion

In this study we report that STING is partly found in the nucleus of various preclinical models and clinical specimens of cytotoxic treatment-resistant breast cancer, and preferentially localizes at the INM. STING promotes the DDR and enhances cancer cell survival in both basal conditions and genotoxic-induced stress. Importantly, these effects are cell-autonomous and independent of the classical CDN/TBK1/IFN inflammatory response, thus identifying a novel functional pathway for STING. Pioneering observations suggest that STING may cooperate with the DDR regulator DNA-PK. Although future work is needed to decipher this new mechanism, the involvement of STING in such a fundamental cellular process as DDR adds a level of complexity to our understanding of this multi-faceted protein in general, and especially in the context of cancer.

STING has been mostly characterized as a transmembrane protein that resides in various cytoplasmic organelles, in agreement with its canonical adaptor function to trigger inflammatory responses upon cytosolic DNA sensing by cGAS (Barber 2015). In this study, we showed by cell fractionation that a part of the STING pool intrinsically resides in the nucleus of various malignant and non-malignant cells, generalizing our preliminary data involving MCF7 cells (Gaston et al. 2016). Its localization to the INM was supported by STING/lamin co-immunostaining and by immunoelectron microscopy analyses. Importantly, STING/lamin co-localization was also observed in PDXs and in all clinical specimens that we analyzed. In their pioneering work, Schirmer and colleagues identified STING as a NET protein (NET23) and tentatively localized it to the ONM using high resolution fluorescence imaging (Schirmer et al. 2003, Malik et al. 2014). INM *versus* ONM localizations may not be mutually exclusive as protein addressing to the INM has been proposed to involve initial protein insertion into ER membranes followed by diffusion to the contiguous ONM and INM (reviewed in Ref. Katta, Smoyer, and Jaspersen 2014). As observed for typical INM-resident proteins such as Emerin (Sullivan et al. 1999), Schirmer team showed that STING failed to localize to the nuclear envelope in lamin A/C-deficient cells (Malik et al. 2010). In addition, STING was shown to participate in chromatin compaction in various cell types, and interaction with epigenetic silencing factors was proposed to occur at the INM (Malik et al. 2014).

The mechanism of STING recruitment to the INM remains to be elucidated. Mechanisms regulating protein addressing to the INM are poorly understood. One model of protein recruitment to the INM suggests a passive diffusion of proteins from ONM to INM through the nuclear pore complexes and retention at the INM via interactions with lamins, chromatin and/or other proteins (Wu, Lin, and Worman 2002, Katta, Smoyer, and Jaspersen 2014). Interestingly, nuclear STING interactome comprises the INM protein TMEM43 that is thought to be involved in nuclear envelope organization, as exemplified by its critical role in Emerin retention at the INM (Bengtsson and Otto 2008). An alternative model proposes that INM proteins contain a nuclear localization signal (NLS)-like motif which is recognized by a karyopherin for an active transport (King, Lusk, and Blobel 2006). Using a NLS prediction software (Kosugi et al. 2009), we found that STING harbors a predicted bipartite NLS motif at the N-terminus (aa 14-46) compatible with both nuclear and cytoplasmic localization. The C-terminally truncated STING-DN isoform contains one additional putative NLS (aa 272-281) exhibiting high confidence score, which is consistent with the preferential accumulation of this isoform in the nuclear fraction that we observed in cell fractionation assays.

This is the first study reporting the involvement of STING in the DDR. STING LOF reduced DDR foci formation (53BP1 and γH2AX) and increased the accumulation of spontaneous and genotoxic-induced DNA breaks. STING GOF had the opposite effect. Accordingly, the proteomic analysis performed in this study revealed that nuclear STING interactome contains several proteins involved in DSB DNA repair: SMARCA5 has been shown to promote DSB repair *via* both NHEJ and HR pathways in an ubiquitin-dependent manner (Smeenk et al. 2013); RUVBL1, RUVBL2 and ACTL6A belong to a histone acetyltransferase complex that modulate 53BP1 DNA binding and the NHEJ/HR balance (Lans, Marteijn, and Vermeulen 2012, Jacquet et al. 2016, Clarke et al. 2017); SUPT16H cooperates with the ubiquitin ligase RNF20 to promote HR (Oliveira et al. 2014); SMC3, a constituent of the cohesion complex, is thought to promote HR by maintaining chromatid sister in close proximity (Potts, Porteus, and Yu 2006). Besides these partners, the most remarkable finding was that nuclear STING interacts with the DNA-PK complex. Although the functional pleiotropy of DNA-PK is emerging (Goodwin and Knudsen 2014, Mohiuddin and Kang 2019), it is mostly known for its role in NHEJ activation and initiation (Neal and Meek 2011). As we showed that STING GOF enhanced DNA-PK protein complex assembly on the chromatin (without affecting their level of expression), it is possible that STING could help recruit and/or stabilize the complex at DNA damage sites.

Interestingly, DNA-PK and STING have already been shown to cooperate in the context of innate immune response to cytosolic DNA. Indeed, co-immunoprecipitation assays from whole-cell extracts revealed the interaction of STING with Ku70 (Ferguson et al. 2012, Morchikh et al. 2017) and DNA-PKcs (Morchikh et al. 2017), and DNA-PK was identified as a DNA sensor triggering inflammatory responses in a STING/TBK1/IRF3-dependent manner (Ferguson et al. 2012, Morchikh et al. 2017). Together with the findings reported in our study, these data establish DNA-PK as a novel canonical STING interactor.

The localization of STING at the INM, where it colocalizes with peripheral DDR foci, is also of particular interest as there is emerging evidence that the nuclear envelope is involved in DNA repair compartmentalization (Marnef and Legube 2017). In yeast (Oza et al. 2009) and drosophila (Ryu et al. 2015), persistent or hard-to-repair DSBs relocate to the nuclear periphery where they are anchored to nuclear pore or INM proteins to be repaired. While DSB mobility to the nuclear periphery has not been described in mammals yet, subnuclear structures with dedicated types of DNA repair have been observed. Hence, in yeast and mammals, DSBs in chromatin domains associated with the nuclear envelope and nuclear pores are preferentially repaired by error-prone pathways, such as NHEJ, alt-NHEJ and Break Induced Replication (BIR) (Lemaitre et al. 2014, Chung et al. 2015). Further studies are required to determine precisely by which mechanism and at which level (i.e. detection, signaling, compartmentalization and/or repair) STING functionally impacts the DDR.

Beyond DNA repair proteins, nuclear STING interactome was also enriched in proteins involved in two other functional networks, namely protein synthesis and mRNA splicing. Interestingly, STING has been shown recently to inhibit host and viral protein synthesis in the context of RNA virus infection. The mechanism was not fully elucidated but involves STING-dependent collapse of polysomes (Franz et al. 2018). Moreover, STING interactor DNA-PK was recently shown to promote ribosome biogenesis, thus impacting global protein synthesis. Indeed, DNA-PK was shown to bind U3 snoRNA to promote pre-rRNA splicing into mature 18S rRNA (Shao et al. 2020). Further work is needed to determine the potential role of STING in those pathways and whether this could impact the DDR.

Based on the evidence that STING is critical for antitumor immune responses, the current clinical trend is to boost STING signaling using STING agonists to enhance tumor eradication by the immune system of the host (reviewed in Ref. Rivera Vargas, Benoit-Lizon, and Apetoh 2017). However, the overall benefit of this therapeutic strategy remains elusive, as recent studies have demonstrated that STING-mediated inflammation can result in pro-tumoral and pro-metastatic effects (Ahn et al. 2014, Gaston et al. 2016, Liu et al. 2018, Bakhoum et al. 2018). Our study identifies DDR as an additional cell-autonomous mechanism by which STING may contribute to tumor progression as well as to resistance to DNA-damaging therapies. Accordingly, in samples of chemotherapy-resistant breast tumors, STING levels (and co-localization with lamina) were particularly elevated in proliferating cells that drive tumor regrowth. This is consistent with clonogenic *in vitro* assays showing that higher STING expression intrinsically promoted survival and regrowth of breast cancer cells exposed to genotoxic stress, while STING deficiency sensitized cells to treatment.

In conclusion, we uncovered a new subcellular localization of STING at the INM of breast cancer cells and provided unprecedented evidence supporting its involvement in the DDR. This newly-identified function highlights a cell-autonomous pathway by which STING promotes cancer cells survival and resistance to DNA-damaging agents. Importantly, the effects of STING on the DDR, cell survival and drug resistance were independent of the canonical CDN/TBK1/IFN pathway. This suggests that in the clinical setting, therapeutic strategies that aim at stimulating canonical STING-mediated antitumor immunity should not promote further STING-mediated DNA repair.

## Materials and Methods

### Cell culture

MCF7 (Estrogen Receptor-positive (ER+); Sigma Aldrich, Saint-Louis, Missouri), BT20 and HCC1937 (triple negative (TN); ATCC, Manassas, Virginie) breast cancer cell lines were purchased between 2008 and 2012. Parental (HEK-Blue™ ISG) and STING-KO (HEK-Blue™ ISG-KO-STING, Invivogen, Toulouse, France) human embryonic kidney (HEK) 293 cells were purchased in 2017. All cell lines were frozen shortly after initial expansion (3-6 passages), and thawed cells were used until passage ∼20. MCF7 and BT20 were authenticated for the last time in December 2019 (STR method). Mycoplasma was tested by PCR once to twice a year. Cells were maintained in DMEM/F12 supplemented with 10% heat-inactivated FBS and 1% penicillin-streptomycin (P/S). Human breast cancer PDX-derived cell lines (HBCx-3 and HBCx-19, ER+; HBCx-2 and HBCx-39, TN) were generated from PDXs developed under IRB approval as previously described (Gaston et al. 2016).

### Patient-derived xenografts (PDX) samples

Paraffin-embedded breast cancer PDX samples (n=4) fixed in 10% neutral buffered formalin were retrieved from the archives of Xentech, Evry, France. Breast cancer PDX establishments and care and use of animals were performed as previously described (Legrier et al. 2016, Marangoni et al. 2007) after approval of the Ethics Committees of the Institut Curie and CEEA-Ile de France Paris (official registration number 59).

### Patient samples

Formalin-fixed, paraffin-embedded breast cancer samples (n=6) were retrieved from the archives of the Department of Pathology, Centre Jean Perrin, Clermont-Ferrand, France. For this study, only samples of breast tumors resistant to neoadjuvant chemotherapy (Fluorouracil-Epirubicin, Cyclophosphamide (FEC)-Taxane regimen), larger than 2 cm in diameter, were used (examples of very limited/partial response to therapy). This study was approved by the Ethics Committee (CECIC) of the Rhone-Alpes-Auvergne region (Grenoble, France).

### Plasmids

pUNO1 (mock), pUNO1-hSTING (STING), pUNO1-hSTING-HA3x (STING-HA) and pUNO1-hSTING-MRP (STING-DN) expression plasmids were purchased from Invivogen. STING-MRP (MITA-related protein) is a spliced variant of STING lacking exon 7 that act as a dominant negative (DN) of STING when it comes to IFN induction due to the lack of binding domains to downstream effectors TBK1 and IRF-3 (1). HA-STING, Flag-STING, Flag-STING-HA and Flag-DN-STING plasmids were generated by subcloning. Plasmids were transfected into cells using Lipofectamine (ThermoFisher, Waltham, Massachussetts). Plasmids psPAX2 and pMD2.G, used for lentiviral particles production, were purchased from Addgene, Watertown, Massachussetts. Lentiviral plasmids containing shRNAs targeting STING (shSTING-1 #TRCN0000161345, shSTING-2 #TRCN0000163029) or non-targeted (scrambled) shRNA (shNT #SHC016-1EA) were purchased from Sigma Aldrich. STING-IRES-GFP and DN-IRES-GFP lentiviral plasmids were generated by inserting the cDNA of STING or STING-DN, respectively, into pWPI backbone (kindly given by M. Orgunc, IUH institute). Empty pWPI lentiviral plasmid was used as a negative control (mock-IRES-GFP).

### Antibodies

We used the anti-STING antibody from Cell Signaling, Danvers, Massachusetts (clone D2P2F, #13647) for the detection of full-length STING, and the anti-STING antibody from R&D Systems, Minneapolis, Minnesota (clone 723505, #MAB7169) only for detection of C-terminally truncated STING-DN by immunoblot.

The antibodies used in this study and their dilution for each experimental procedure (IF, immunoflouorescence; IHF: immunohistofluorescence; IP, immunoprecipitation) are listed in Table 1.

**Table 1:**
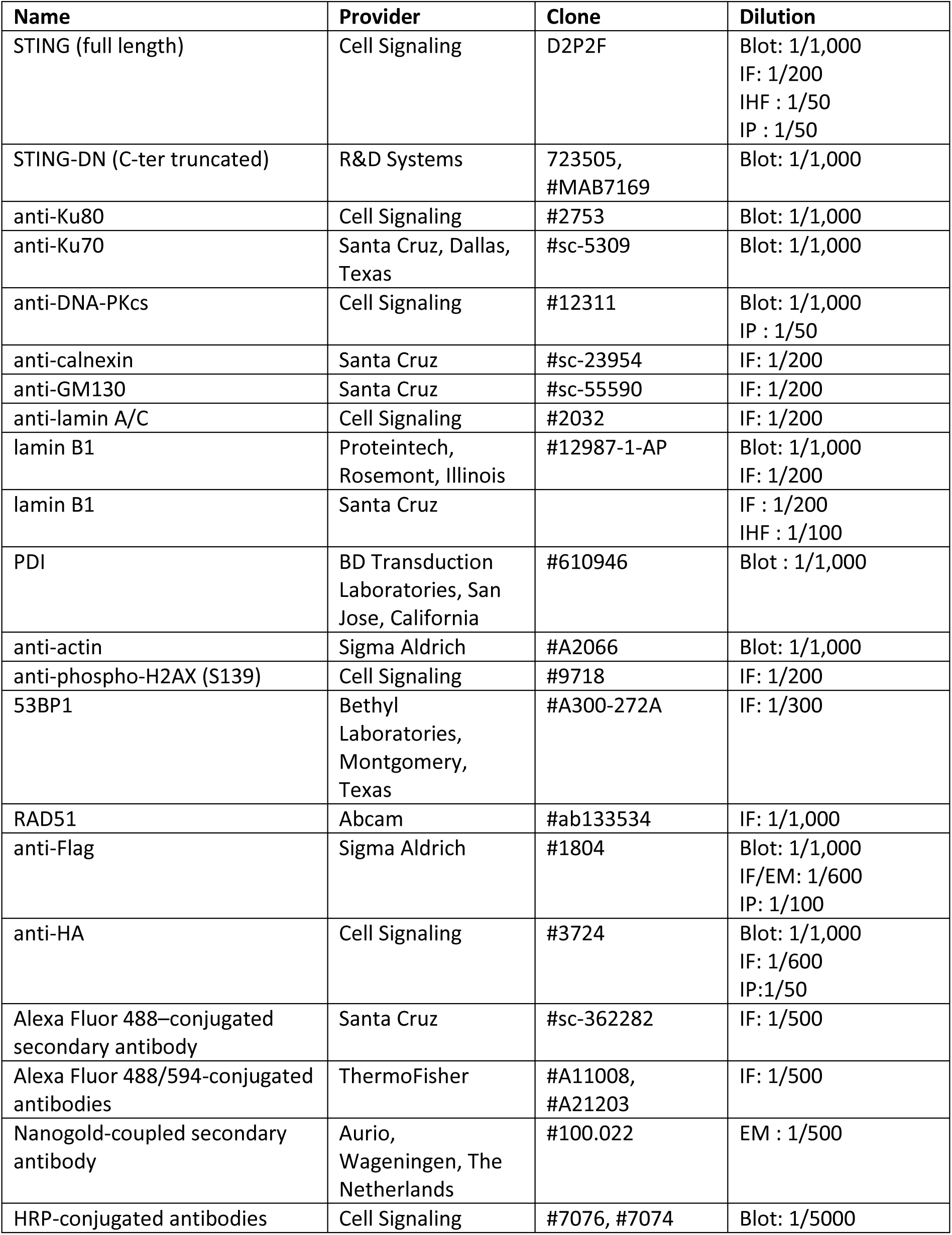
list of antibodies

| **Name** | **Provider** | **Clone** | **Dilution** |
| --- | --- | --- | --- |
| STING (full length) | Cell Signaling | D2P2F | Blot: 1/1,000<br>IF: 1/200<br>IHF : 1/50<br>IP : 1/50 |
| STING-DN (C-ter truncated) | R&D Systems | 723505,<br>#MAB7169 | Blot: 1/1,000 |
| anti-Ku80 | Cell Signaling | #2753 | Blot: 1/1,000 |
| anti-Ku70 | Santa Cruz, Dallas,<br>Texas | #sc-5309 | Blot: 1/1,000 |
| anti-DNA-PKcs | Cell Signaling | #12311 | Blot: 1/1,000<br>IP : 1/50 |
| anti-calnexin | Santa Cruz | #sc-23954 | IF: 1/200 |
| anti-GM130 | Santa Cruz | #sc-55590 | IF: 1/200 |
| anti-lamin A/C | Cell Signaling | #2032 | IF: 1/200 |
| lamin B1 | Proteintech,<br>Rosemont, Illinois | #12987-1-AP | Blot: 1/1,000<br>IF: 1/200 |
| lamin B1 | Santa Cruz |  | IF : 1/200<br>IHF : 1/100 |
| PDI | BD Transduction<br>Laboratories, San<br>Jose, California | #610946 | Blot : 1/1,000 |
| anti-actin | Sigma Aldrich | #A2066 | Blot: 1/1,000 |
| anti-phospho-H2AX (S139) | Cell Signaling | #9718 | IF: 1/200 |
| 53BP1 | Bethyl<br>Laboratories,<br>Montgomery,<br>Texas | #A300-272A | IF: 1/300 |
| RAD51 | Abcam | #ab133534 | IF: 1/1,000 |
| anti-Flag | Sigma Aldrich | #1804 | Blot: 1/1,000<br>IF/EM: 1/600<br>IP: 1/100 |
| anti-HA | Cell Signaling | #3724 | Blot: 1/1,000<br>IF: 1/600<br>IP:1/50 |
| Alexa Fluor 488–conjugated<br>secondary antibody | Santa Cruz | #sc-362282 | IF: 1/500 |
| Alexa Fluor 488/594-conjugated<br>antibodies | ThermoFisher | #A11008,<br>#A21203 | IF: 1/500 |
| Nanogold-coupled secondary<br>antibody | Aurio,<br>Wageningen, The<br>Netherlands | #100.022 | EM : 1/500 |
| HRP-conjugated antibodies | Cell Signaling | #7076, #7074 | Blot: 1/5000 |

### Generation of stable cell lines

MCF7 cells stably overexpressing Flag and/or HA-tagged isoforms of STING were obtained by blasticidin selection (100 µg/mL) of cells transfected with cognate plasmids. The knockdown and rescue of STING expression in MCF7 cells were performed using the lentivirus technology. Briefly, HEK293T cells (a kind gift from Simon Fillatreau, Inserm U1151) were transfected in antibiotic-free DMEM medium with psPAX2, pMD2.G and shSTING-1, shSTING-2 or shNT plasmids. Culture medium was replaced by fresh DMEM/F12 medium the day after transfection. Supernatant containing viral particles were collected 48h after transfection, centrifuged (500 g, 5 min), filtered (PES 45µm) then added onto MCF7 cells. The day after transduction, shSTING-1, shSTING-2 and shNT cells were selected with puromycin (2 µg/mL) for one week. The same procedure was used to generate the rescued cell lines: shNT cells were transduced with viruses containing mock-IRES-GFP plasmid (“shNT + mock” cells) while shSTING-2 cells were transduced with viruses containing either mock-IRES-GFP (shSTING + mock), STING-IRES-GFP (shSTING + STING) or STING-DN-IRES-GFP (shSTING + STING-DN) plasmid. Stably transduced cells were GFP-sorted by flow cytometry.

### Transfection of siRNAs

Using interferin reagent (Polyplus transfection, Illkirch-Graffenstaden, France), cells were transfected 24h post-seeding with the following siRNA (GE Dharmacon, Lafayette, Colorado): siNonTargeted (D-001810-10), siSTING (L-024333-02), siIFNAR1 (L-020209-00) and siTBK1 (L-003788-00). For siRNA efficiency, cells were analyzed 3 days post-transfection. For other experiments, cells were analyzed at least 2 days after transfection, as indicated.

### RT-qPCR

Total RNA was extracted using the NucleoSpin RNA XS kit (Macherey-Nagel, Gutenberg, France). One microgram of total RNA was reverse-transcribed into cDNA using High Capacity cDNA Reverse Transcription Kit (Applied biosystems, Foster City, California). Gene expression was analyzed with SYBR Select Master Mix (Life Technologies, Carlsbad, California). RT-qPCR data were normalized to the expression levels of housekeeping genes: GAPDH (glyceraldehyde 3-phosphate dehydrogenase) and RPL13 (ribosomal protein L13).

### Cell fractionation

Sequential fractionation was adapted from a previous report (Baghirova et al. 2015). Cells were lysed (30 min, 4°C) in cytoplasm-extraction buffer (50 mM Hepes, pH 7.4, 150 mM NaCl, 1% v:v NP-40) then centrifuged 5 min at 7,000 g and the supernatant (cytoplasm fraction, C) was collected. After 3 washes in same buffer, nucleus-containing pellets were resuspended and incubated (1h, 4°C) in nucleus-extraction buffer (RIPA, 2 mM MgCl2 and 50 U/mL benzonase). Lysates were centrifuged 5 min at 2,000 g to remove remaining insoluble cellular debris and the supernatant (nucleus fraction, N) was recovered.

### Chromatin fraction

Chromatin fractionation was performed as previously described with minor modifications (Ochi et al. 2015). Cells were pre-extracted twice (3 min on ice) with CSK buffer (10mM Hepes, 100 mM NaCl, 300 mM sucrose, 3 mM MgCl2) containing 0.7% Triton X-100 and 0.3 mg/mL RNase A. Next, cells were washed twice in ice-cold PBS then lysed (1h, 4°C) in nucleus-extraction buffer (see above). The chromatin fraction was recovered in the supernatant after centrifugation (5 min at 2,000 g). For whole cell lysates, the pre-extraction step was skipped.

### Co-immunoprecipitation

Nucleus-containing pellets (see Cell fractionation protocol) were lysed (1h, 4°C) in low-denaturing lysis buffer (50 mM Hepes pH 7.4, 50 mM Tris-HCl pH 7.4, 150 mM NaCl, 1 mM EDTA, 1% v:v Triton X100) supplemented with 2 mM MgCl2 and 50 U/mL benzonase. Nuclear lysates were incubated overnight at 4°C with/without the indicated antibody then immune complexes were captured by addition of protein A/G magnetic beads (ThermoFisher) for 2h at RT. After 3 washes in low-denaturing lysis buffer, immune-complexes were denatured in LDS (4X)/β-mercaptoethanol (20%) sample buffer.

### Immunoblotting

Cell lysate protein concentration was determined using a Micro BCA Protein Assay kit (Bio Basic). Unless specified, proteins were diluted in LDS (1X)/β-mercaptoethanol (5%) sample buffer, denaturated 5 min at 95°C and finally loaded on NuPAGE 4-12% Bis-Tris protein gels (Life Technologies) and electro-transferred onto nitrocellulose membranes. Membranes were blocked (3% BSA, 30 min, RT), then incubated with primary (overnight, 4°) and secondary (1h, RT) antibodies. Membranes were revealed with suitable HRP substrates (Clarity ECL Western Blotting Substrate, Biorad, Hercumes, California and Immobilion ECL Ultra Western HRP Substrate, Merck, Darmstadt, Allemagne) and quantified by Image Lab software (Biorad)

### NanoLC-MS/MS protein identification and quantification

S-TrapTM micro spin column (Protifi, Huntington, New York) digestion was performed on IP eluates according to manufacturer’s protocol. Samples were digested with 2µg of trypsin (Promega, Madison, Wisconsin) at 37°C overnight. After elution, peptides were finally vacuum dried down. Samples were resuspended in 35 µL of 10% ACN, 0.1% TFA in HPLC-grade water. For each run, 5 µL was injected in a nanoRSLC-Q Exactive PLUS (RSLC Ultimate 3000) (ThermoFisher). Peptides were loaded onto a µ-precolumn (Acclaim PepMap 100 C18, cartridge, 300 µm i.d.×5 mm, 5 µm) (ThermoFisher), and were separated on a 50 cm reversed-phase liquid chromatographic column (0.075 mm ID, Acclaim PepMap 100, C18, 2 µm) (ThermoFisher). Chromatography solvents were (A) 0.1% formic acid in water, and (B) 80% acetonitrile, 0.08% formic acid. Peptides were eluted from the column with the following gradient 5% to 40% B (120 minutes), 40% to 80% (1 minute). At 121 minutes, the gradient stayed at 80% for 5 minutes and, at 127 minutes, it returned to 5% to re-equilibrate the column for 20 minutes before the next injection. One blank was run between each series to prevent sample carryover. Peptides eluting from the column were analyzed by data dependent MS/MS, using top-10 acquisition method. Peptides were fragmented using higher-energy collisional dissociation (HCD). Briefly, the instrument settings were as follows: resolution was set to 70,000 for MS scans and 17,500 for the data dependent MS/MS scans in order to increase speed. The MS AGC target was set to 3.106 counts with maximum injection time set to 200 ms, while MS/MS AGC target was set to 1.105 with maximum injection time set to 120 ms. The MS scan range was from 400 to 2000 m/z. Dynamic exclusion was set to 30 seconds duration.

### Data Processing Following LC-MS/MS acquisition

The MS files were processed with the MaxQuant software version 1.5.8.3 and searched with Andromeda search engine against the database of Homo Sapiens from swissprot 07/2017. To search parent mass and fragment ions, we set an initial mass deviation of 4.5 ppm and 20 ppm respectively. The minimum peptide length was set to 7 aminoacids and strict specificity for trypsin cleavage was required, allowing up to two missed cleavage sites. Carbamidomethylation (Cys) was set as fixed modification, whereas oxidation (Met) and N-term acetylation were set as variable modifications. Match between runs was not allowed. LFQ minimum ratio count was set to 1. The false discovery rates (FDRs) at the protein and peptide level were set to 1%. Scores were calculated in MaxQuant as described previously (Cox J., Mann M., 2008). The reverse and common contaminants hits were removed from MaxQuant output. Proteins were quantified according to the MaxQuant label-free algorithm using LFQ intensities [Luber, 2010 #1907; Cox, 2008 #1906]. Samples were analysed in triplicates and data were analyzed with Perseus software (version 1.6.2.3) freely available at www.perseus-framework.org. The LFQ (Label-free Quantification) data were transformed in log2. All the proteins identified in all of the 3 replicates were submitted to statistical test (volcano plot, FDR=0.001 and S0=0.5) after imputation of the missing value by a Gaussian distribution of random numbers using default settings. Protein annotations (GO, Keywords) were retrieved directly using via perseus.

### Immunofluorescence

For immunohistofluorescence experiments, formalin-fixed paraffin-embedded samples underwent deparaffination and rehydration followed by heat-induced antigen retrieval in citrate buffer (pH 6). For immunocytofluorescence experiments, cells were grown in sterile chamber slides (#80826, IBIDI, Gräfelfing, Germany), treated as indicated, then fixed in 4% paraformaldehyde and permeabilized in 0.1% Triton X100. Both types of samples were then blocked with 3% Bovine Serum Albumin/2% normal goat serum before overnight incubation at 4°C with primary antibodies and 1h incubation at RT with fluorescent secondary antibodies. Nuclei were stained with DAPI (1µg/mL) for 10 min at RT (ThermoFisher). Acquisition was performed using an Apotome Zeiss at 10X (comet assays), 20X (immunohistofluroescence and 53BP1 foci) and 63X (co-localization assays) magnification. Merged images were treated by Image J software to emphasize co-localization (appearing in white).

### Electron microscopy

Cells collected in 1,5 mL Eppendorf tube were fixed, permeabilized and blocked as for immunofluorescence analyses. Cells were incubated overnight (4°C) with/without anti-Flag antibody then incubated (1h at RT) with nanogold-coupled secondary antibody and finally fixed with 2.5% glutaraldehyde EM grade (Sigma Aldrich, #16210) for 1 h. Sample were post-fixed with 1% osmium tetroxide (EMS) in 0.1 M phosphate buffer then gradually dehydrated in 70, 90 and 100% ethanol. After 10 min in a 1:2 mixture of epoxy propane and epoxy resin and 10 min in epon, samples were embedded in epoxy resin and polymerized at 60°C for 24 h. After polymerization, ultrathin sections (90 nm) were cut with an ultra-microtome (Reichert ultracut S) on 100 mesh grids (Gilder), stained with uranyl acetate and Reynold’s lead and observed with a transmission electron microscope (JEOL 1011). Acquisition was performed with a Gatan Orius 1000 CCD camera.

### Comet assay

MCF7 cells were treated with mafosfamide (10µM) one day post-seeding and maintained for another 6 days which corresponds to maximal γH2AX accumulation (Gaston et al. 2016). When relevant, cells were transfected with expression plasmids 8h prior treatment. Naïve HEK293 cells were transfected one day post-seeding with either mock or STING plasmids (as indicated) and maintained for 3 days. Comet assays were performed according to the manufacturer’s instructions (4250-050-K; Trevigen, Gaithersburg, Maryland). DNA damage was measured in terms of tail moments using OpenComet plugin in ImageJ software.

### Cell viability assay

Cells were treated with vehicle, mafosfamide or etoposide (10µM or as indicated) one day post-seeding. In the siRNA/genotoxic combination setting, cells were transfected one day post-seeding and treated with mafosfamide 3 days later. Cell viability was measured using CellTiter-Glo Luminescent cell viability assay reagent. Luminescence was measured with a microplate reader (Mithras LB940, Berthold, Bad Wildbad, Germany) and data were normalized to the vehicle and/or non-targeted siRNA/shRNA (siNT/shNT) condition as indicated.

### Colony assay

MCF7 cells were treated with 10µM mafosfamide one day post-seeding. BT20 cells were transfected with siRNAs one day post-seeding and treated with 10µM mafosfamide 3 days later. When indicated, cells were transfected with expression plasmids the day of treatment. After 20 or 40 days, colonies were stained and quantified using the plugin ColonyArea in ImageJ software.

### Statistical analysis

Data were analyzed using GraphPad Prism 6 software (GraphPad, San Diego, California) and are presented as mean +/- s.d. unless otherwise indicated. For comparison between two groups two-tailed Student’s t-test was used while comparisons between multiple groups were performed using one-way ANOVA. To examine the influence of two independent parameters on multiple groups two-way ANOVA was used. Statistically significant differences are indicated as follows: *p < 0.05; **p <0.01; ***p < 0.001; ****p < 0.0001. Each experiment was repeated independently three times unless otherwise indicated. Statistical details of experiments can be found in the figure legends.

## Acknowledgements

This study was funded by XenTech, annual funding from Inserm and the University Paris Descartes and ECLER non-profit association. JG and LC were recipient of a CIFRE fellowship from the Association Nationale de la Recherche et de la Technologie (ANRT). MM is supported by the French National League against Cancer.

The authors would like to thank Nicolas Goudin for helpful assistance with imaging, and the electron microscopy facility of Hôpital Cochin. We are grateful to Patrice Codogno, Etienne Morel, Ganna Panasyuk, Thierry Dubois and Frédéric Rieux-Laucat for helpful discussions, to Dr Orgunc and Pr S. Fillatreau for providing biological materials, and to Katheryn Meek for critical reading of the manuscript.

## Author Contributions

Conception and design: L. Cheradame, J. Gaston, S. Cairo, V. Goffin

Development of methodology: L. Cheradame, I. C. Guerrera, A. Schmitt, S. Cairo, V. Goffin

Acquisition of data: L. Cheradame, I. C. Guerrera, A. Schmitt, V. Jung, M. Pouillard

Analysis and interpretation of data: L. Cheradame, I. C. Guerrera, A. Schmitt, N. Radosevic-Robin, M. Modesti, S. Cairo, V. Goffin

Writing, review, and/or revision of the manuscript: L. Cheradame, I. C. Guerrera, A. Schmitt, M. Modesti, J.-G. Judde, S. Cairo, V. Goffin

Administrative, technical, or material support: L. Cheradame, I. C. Guerrera, A. Schmitt, V. Jung, N. Radosevic-Robin, J.-G. Judde, S. Cairo, V. Goffin

Study supervision: S. Cairo, V. Goffin

## Supplemental Information

**Fig S1.**
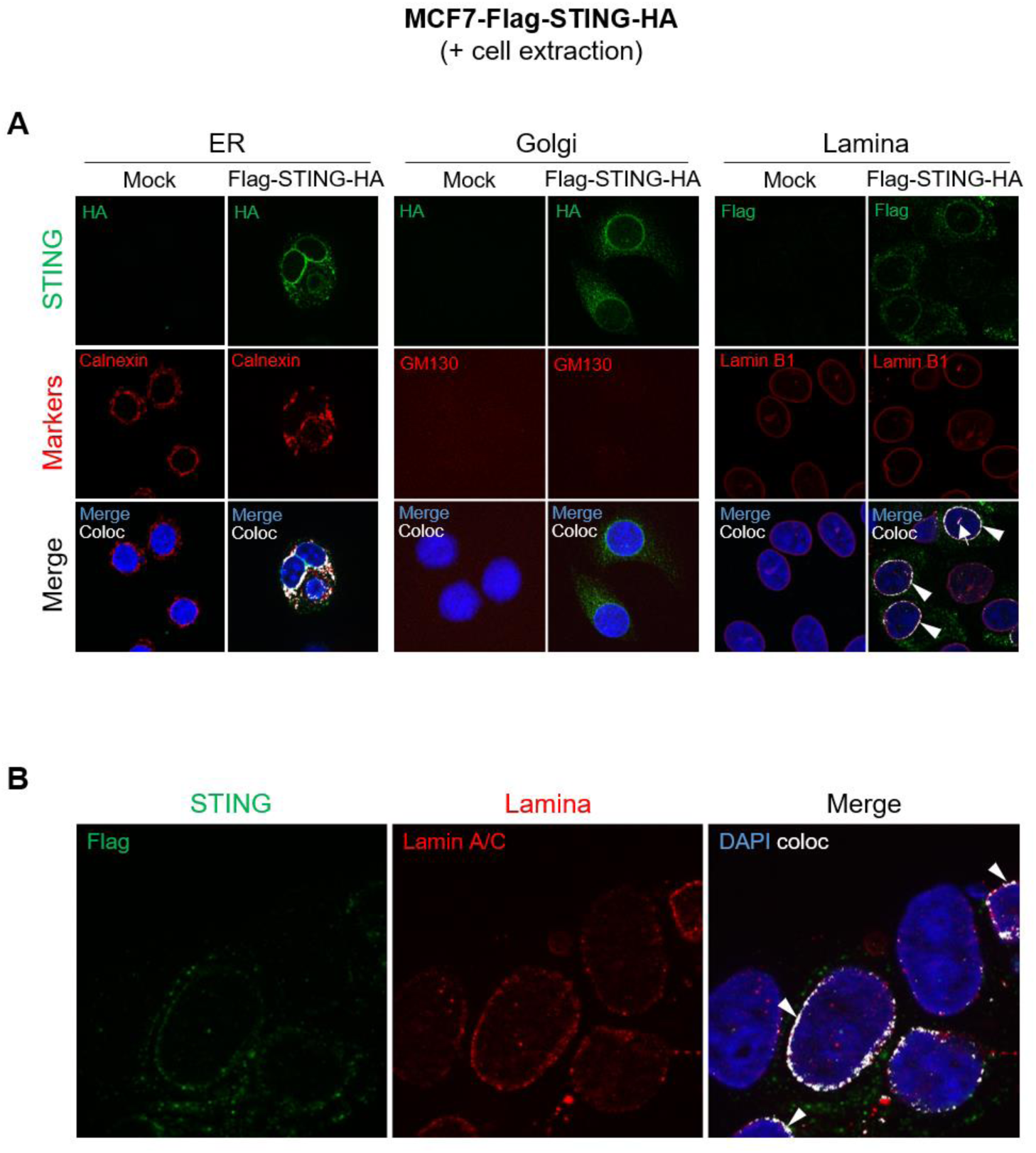
Nuclear STING co-localizes with the lamina in breast cancer cells. **a**,**b** The same experiment as in main Fig2a was performed after pre-extraction treatment of MCF7 cells ectopically expressing Flag-STING-HA *versus* empty vector (Mock). STING was stained using anti-HA or anti-Flag antibodies (according to the species of the antibody directed against subcellular markers) as indicated. Middle panels show immunofluorescence of ER (calnexin), Golgi (GM130) and nuclear lamina (lamin B1) markers in **a**, and of lamin A/C in **b**. Lower (**a**) and right (**b**) panels display merged images. Nuclei were stained with DAPI. White arrowheads point to co-localization of STING at the lamina rim, and white arrows to intra-nuclear staining.

**Fig S2.**
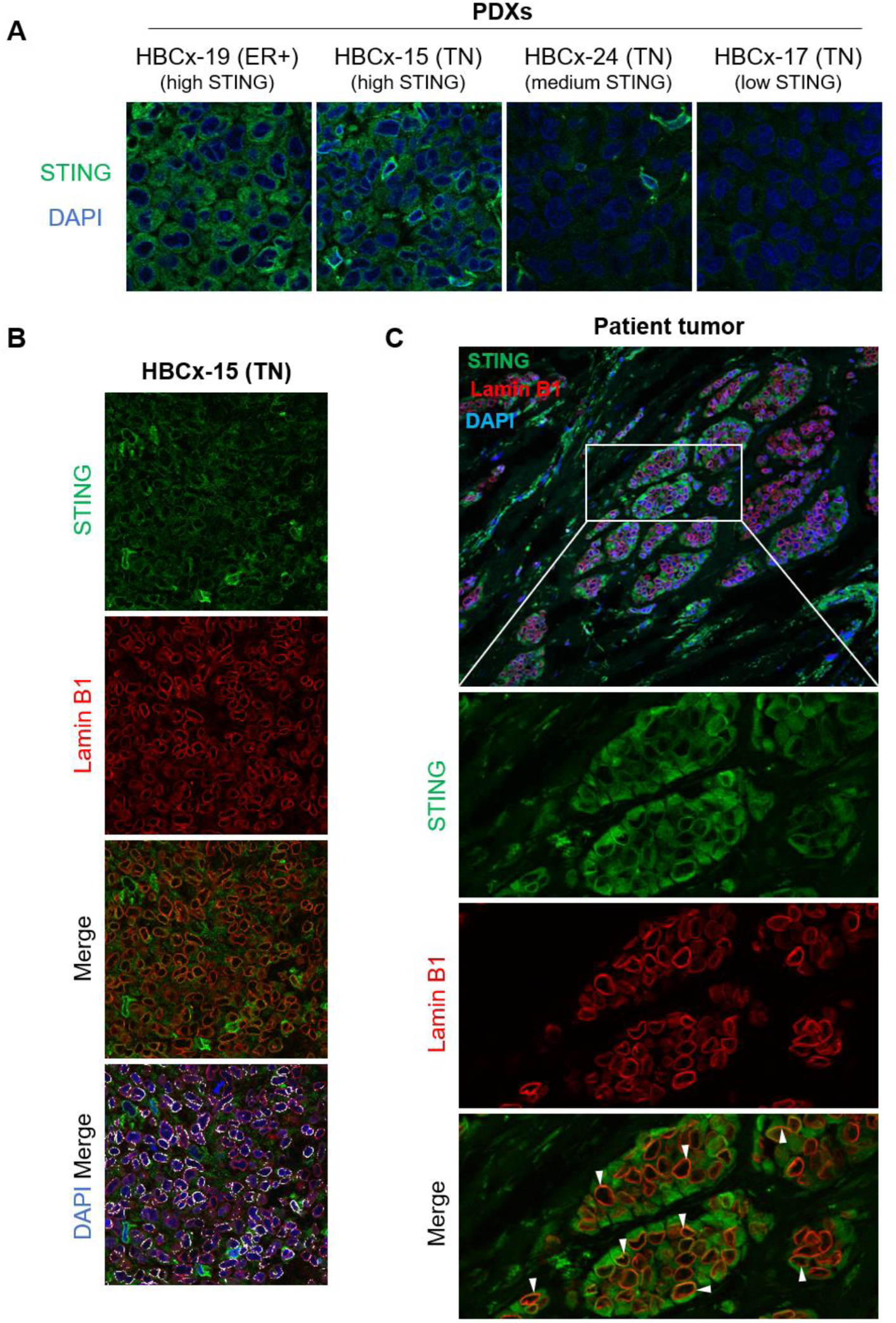
Nuclear STING co-localizes with the lamina in breast cancer cells PDXs and patient breast tumors. **a** Examples of endogenous STING immunostaining in ER+ and TN breast cancer PDXs exhibiting different levels of STING as determined by transcriptomic analysis (high, medium, low). **b** Immunofluorescence of endogenous STING and lamin B1 in the high STING-expressing HBCx-15 PDX. Lower panels display merged images with or without DAPI and image treatment by Image J software to emphasize co-localization (appearing in white). **c** The upper image shows one representative patient tumor area immunostained for STING and lamin B1 (nuclei stained with DAPI). The three bottom panels show higher magnification of the squared area for STING, lamin B1 and merged staining, as indicated. White arrowheads: examples of STING/lamin B1 co-localization.

**Fig S3.**
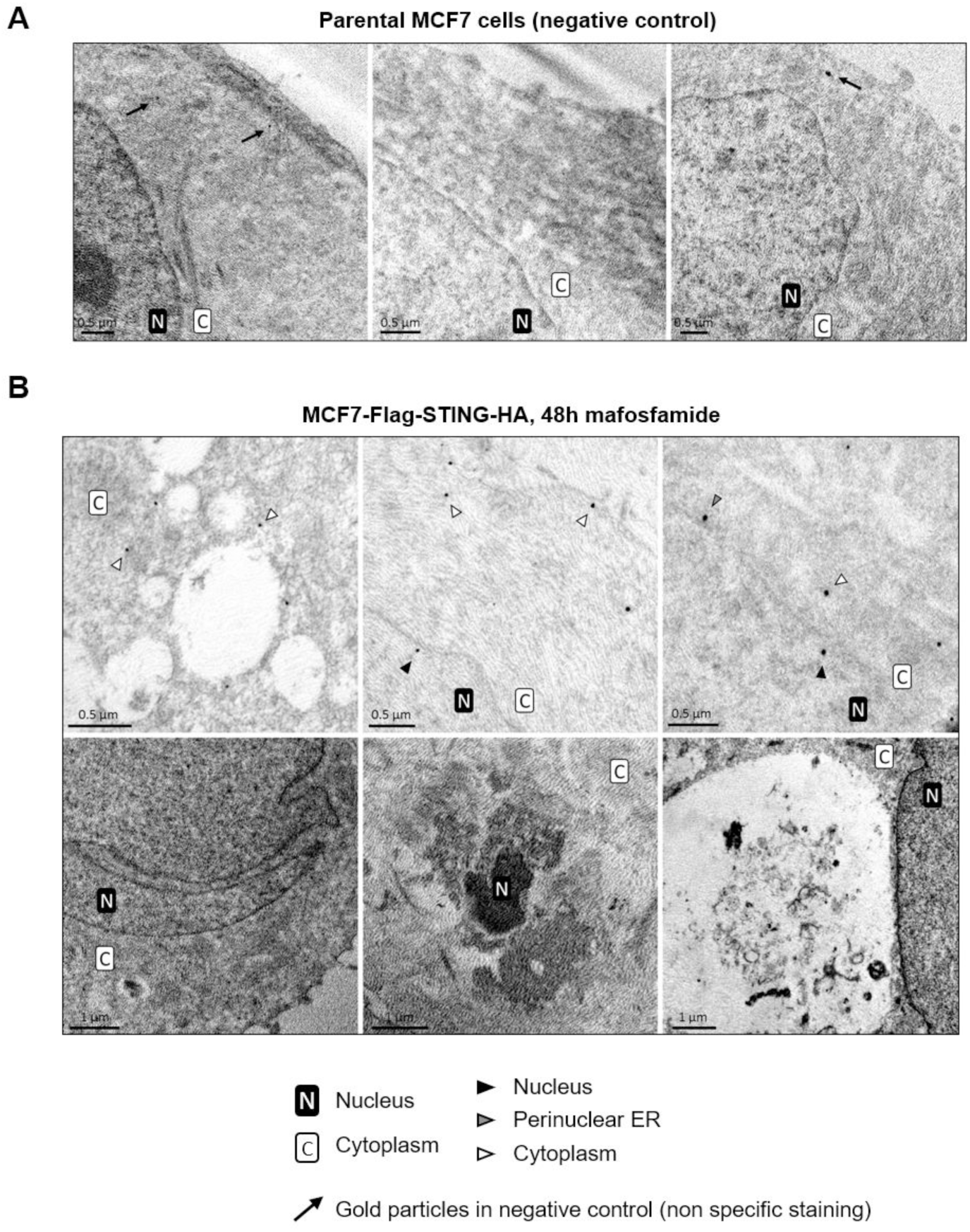
Localization of STING as assessed by immunoelectron microscopy. **a** Representative images of parental MCF7 cells stained with anti-flag immunogold labeling procedure used as the negative control of experiments shown in Fig. 3. Black arrows indicate minimal nonspecific nanogold staining. **b** Six representative images of MCF7 cells expressing Flag-STING-HA after mafosfamide treatment (48h). Upper panels: anti-flag immunogold staining showing STING-positive vesicles (left) and STING present at both sides of the nuclear membrane (middle and right). Lower panels: mafosfamide-treated cells displayed several signs of stress including nuclei with irregular shape and large invaginations, picnotic nuclei, dilated ER with dramatically enlarged lumen and large vesicles filled with cell debris. STING subcellular localization is identified using symbols displayed below the figure. Size bars are indicated in each panel.

**Fig S4.**
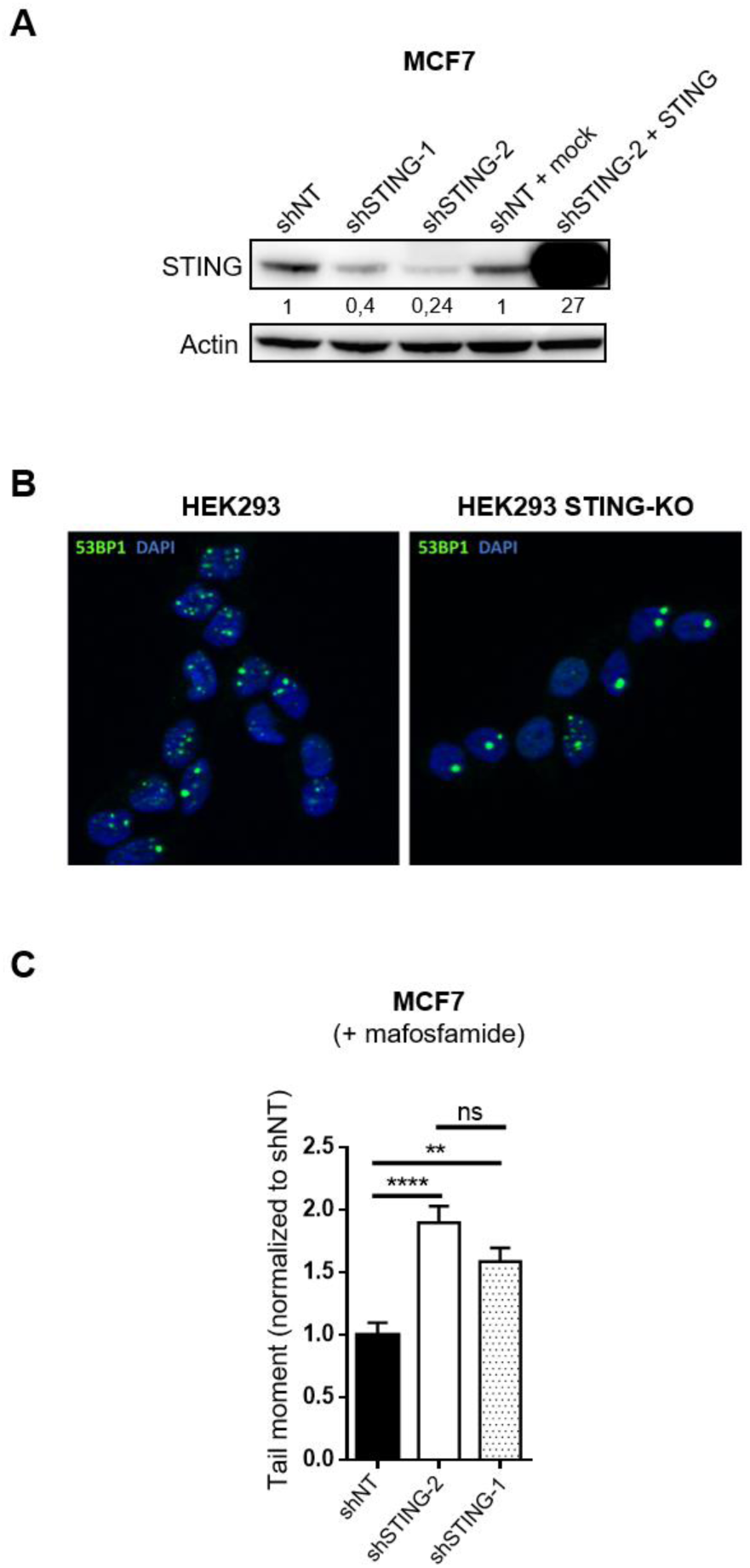
STING contributes to the DDR. **a** Immunoblot of STING in MFC7 cells stably transduced with either non-targeted shRNA (shNT) or two different shRNAs targeting *STING* (shSTING-1 and shSTING-2) and rescued using STING-encoding *versus* empty vectors (mock), as indicated. **b** Immunofluorescence of 53BP1 foci in naïve parental (WT) and STING-deficient (KO) HEK293 cells. Nuclei were stained with DAPI. Quantification is shown in main Fig 4c. **c** Tail moment after mafosfamide treatment of MCF7 cells stably transduced with shNT or two different shRNAs targeting *STING* (shSTING-2 and shSTING-1). Mean ± s.e.m of tail moment of n=746 (shNT), n=976 (shSTING-2) and n= 1,084 (shSTING-1) cells from n=3 independent experiments (one-way ANOVA and post-hoc Tukey’s multipe comparison test).

**Fig S5.**
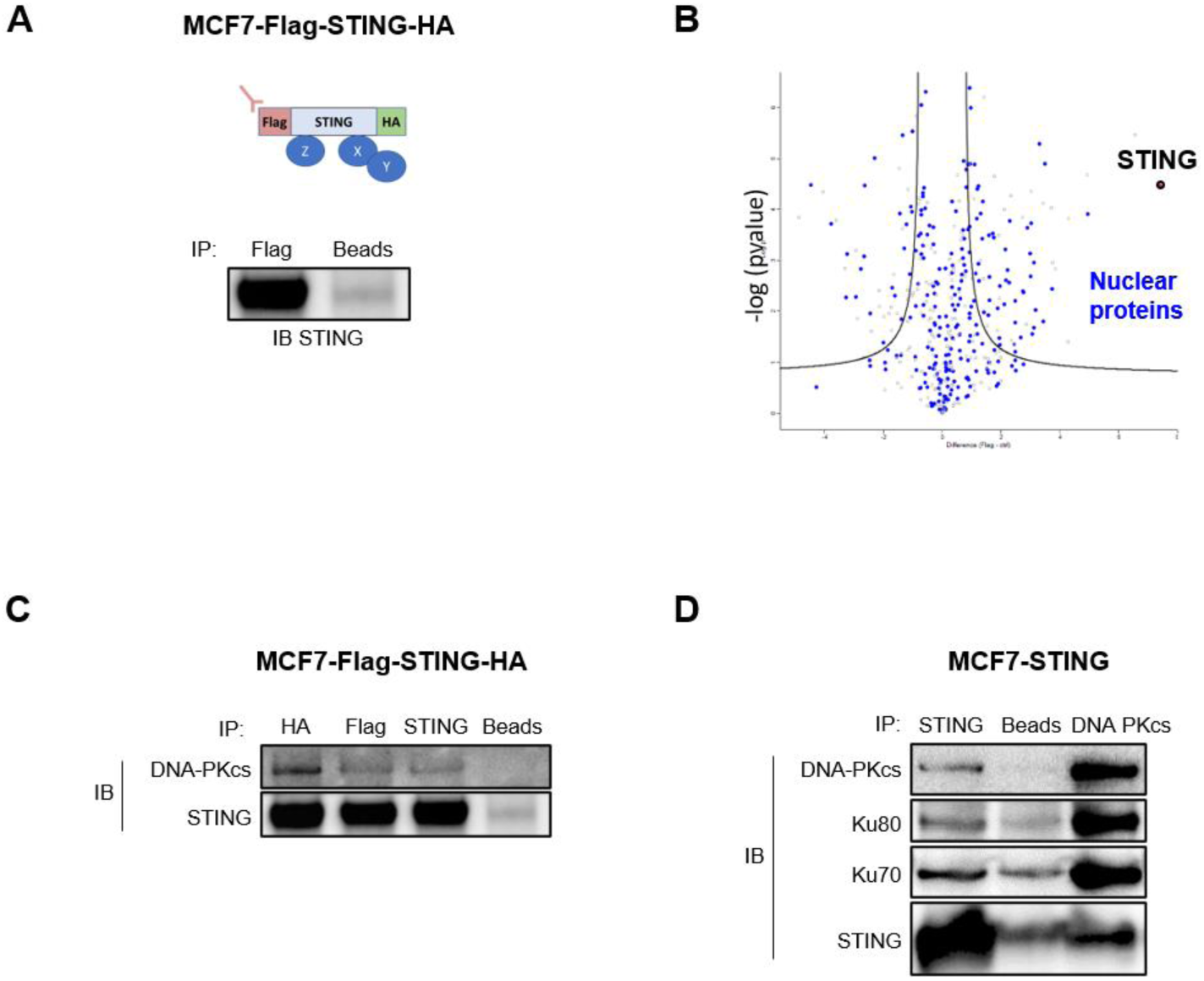
Nuclear STING interactome by mass spectrometry. **a** Immunoblot of STING following Flag-STING-HA immunoprecipitation from nuclear extracts of stably transfected MCF7 cells using anti-Flag antibodies or beads only (negative control). **b** Volcano plot of –log(*p*value) *versus* fold change of proteins present in anti-STING immunoprecipitates *versus* negative control as detected by mass spectrometry. Nuclear proteins (Gene Ontology Cell Component GOCC and Keywords of Uniprot databases) are colored in blue. **c** Immunoblots (IB) of DNA-PKcs in immunoprecipitates recovered using three antibodies mapping distinct regions of Flag-STING-HA ectopically expressed in MCF7 cells. **d** Immunoblots of proteins constituting the DNA-PK complex (DNA-PKcs, Ku80, Ku70) in immunopecipitates of untagged STING ectopically expressed in MCF7 cells. The reverse immunoprecipation was performed using anti-DNA-PKcs antibody (right lane). In **c-d**, the negative control involved beads only.

**Supplementary Table 1** (provided as a separate excel file).

### Protein partners of nuclear STING (TMEM173)

List of protein identified after immunoprecipitation of STING using anti-Flag antibody (Fla1 to 3) and negative control (ctrl1 to 3). For each protein we report the log(2) of the LFQ values. Values in grey are imputed values as described in material and method section, as these proteins were not identified and/or quantified in that samples. The t-test was performed between the “ctrl” and “Flag” after imputation of the missing values. Protein identified with a FDR=0.001, S0=0.5 are marked as +. In green, we highlight the proteins significantly higher in the Flag samples. Proteins classified as “Nucleus” in GOCC and Keywords of Uniprot are reported in light blue (as in Figure S5b). In red, proteins belonging to the DNA repair as highlighted in Figure 7b.

